# Florigen and antiflorigen gene expression correlates with reproductive state in a marine angiosperm, *Zostera marina*

**DOI:** 10.1101/2024.11.09.622789

**Authors:** Christine T. Nolan, Ian Campbell, Anna Farrell-Sherman, Bryan A. Briones Ortiz, Kerry A. Naish, Verónica Di Stilio, James E. Kaldy, Cinde Donoghue, Jennifer L. Ruesink, Takato Imaizumi

## Abstract

- Florigen and antiflorigen genes within the phosphatidylethanolamine-binding protein (PEBP) family regulate flowering in angiosperms. In eelgrass (*Zostera marina*), a marine foundation species threatened by climate change, flowering and seed production are crucial for population resilience. Yet, the molecular mechanism underpinning flowering remains unknown.
- Using phylogenetic analysis and functional assays in *Arabidopsis*, we identified thirteen *PEBP* genes in *Z. marina* (*ZmaPEBP*) and showed that four genes altered flowering phenotypes when overexpressed. We used quantitative RT-PCR on *Z. marina* shoots from perennial and annual populations in Willapa Bay, USA to assess expression of these four genes in different tissue and expression changes throughout the growth season.
- We demonstrated that *ZmaFT2* and *ZmaFT4* promote flowering, and *ZmaFT9* and *ZmaTFL1a* repress flowering in *Arabidopsis*. Across five natural sites exhibiting different degrees of population genetic structure, *ZmaFT2* and *ZmaFT4* were expressed in leaves of vegetative and reproductive shoots and in stems and rhizomes of reproductive shoots. *ZmaFT9* was distinctively expressed in leaves of vegetative and juvenile shoots, while *ZmaTFL1a* levels increased after flowering shoots emerged.
- Our results suggest that *ZmaFT2* and *ZmaFT4* may promote flowering, while *ZmaFT9* may inhibit a floral transition in eelgrass. We speculate that *ZmaTFL1a* may be involved in flowering shoot architecture.

## INTRODUCTION

In angiosperms, flowering is induced by florigen (flowering-inducing substrate) genes, with *FLOWERING LOCUS T* (*FT*) as the main inducer of flowering processes. *FT* is a member of a larger family of genes that encode phosphatidylethanolamine-binding proteins (PEBP), which includes other genes relevant to flowering, all of which are highly conserved across flowering plants (Pin & Nilsson, 2012; Wickland & Hanzawa, 2015). In *Arabidopsis thaliana*, there are six *FT*-like genes implicated in flowering and reproductive processes. *FT* and *TWIN SISTER OF FT* (*TSF*) generate small proteins that move through the plant to the shoot apical meristem, and both activate flowering (florigen); as such, they have redundant effects (Kardailsky *et al*., 1999; Yamaguchi *et al*., 2005; Corbesier *et al*., 2007). *TERMINAL FLOWER 1* (*TFL1*), *CENTRORADIALIS* (*ATC*), and *BROTHER OF FT AND TFL1* (*BFT*) all inhibit the onset of flowering (Kobayashi *et al*., 1999; Mimida *et al*., 2001; Yoo *et al*., 2010). *MOTHER OF FT AND TFL1* (*MFT*), while capable of activating flowering processes, also plays a role in seed germination (Yoo *et al*., 2004; Xi *et al*., 2010). In *Arabidopsis*, *PEBP* genes involved with the photoperiodic flowering pathway show tissue-specific expression patterns. *FT* is primarily expressed in leaf phloem companion cells, generating a protein that acts as a long distance signal transported through the phloem (Takada & Goto, 2003; Corbesier *et al*., 2007). *TFL1* expression occurs in meristem tissue in shoots in *Arabidopsis* and both *FT* and *TFL1* play a role in shoot indeterminacy and maintenance of vegetative state (Bradley *et al*., 1997; Ratcliffe *et al*., 1999; Baumann *et al*., 2015; Liu *et al*., 2023b). The *PEBP* gene family has undergone expansion in monocot lineages, specifically within the *FT* clade, with species like *Oryza sativa* and *Brachypodium distachyon* having 19 and 18 members of their *PEBP* gene families (13 and 12 within *FT* clades), respectively (Itoh *et al*., 2010; Wu *et al*., 2013; Bennett & Dixon, 2021; Liu *et al*., 2023a). *PEBP* genes have multifaceted roles in plant growth and development in other plant species; *FT* and *TFL1* are both known to affect branching and shoot architecture in flowering plants including *Arabidopsis* and tomato (Niwa *et al*., 2013; Lifschitz *et al*., 2014; Weng *et al*., 2016). *FT* homologs have also been shown to promote tuber and stolon formation in potatoes and strawberries, respectively (Navarro *et al*., 2011; Gaston *et al*., 2021). *FT* expression is mainly controlled by photoperiod (daylength) and temperature, among other environmental conditions, which in turn regulates flowering time (Blázquez *et al*., 2003; Balasubramanian *et al*., 2006; Lee *et al*., 2007; de Montaigu *et al*., 2010; Song *et al*., 2013; Kinmonth-Schultz *et al*., 2016; Susila *et al*., 2018; Takagi *et al*., 2023).

Because flowering directly impacts seed production and fruit yield in plant reproduction, *FT* function and the floral pathway have been extensively studied in a wide breadth of terrestrial plant species (Wickland & Hanzawa, 2015), including dicots such as *Beta vulgaris* (Pin *et al*., 2010) and *Chrysanthemum setiscupe* (Oda *et al*., 2012), and also monocot species such as *Allium cepa* (Lee *et al*., 2013) and *Oryza sativa* (Kojima *et al*., 2002). Characterization of *FT* function in aquatic species has only been recently explored in fresh-water species (Yoshida *et al*., 2021), and has not yet been investigated in marine angiosperm species. Eelgrass (*Zostera marina*) is a seagrass native to both the Atlantic and Pacific Northern Hemisphere, one of about 60 species of marine angiosperms and a member of the early-diverging monocot order *Alismatales*. Despite its small genome of 202.3 Mb (Olsen *et al*., 2016), eelgrass is predicted to have at least thirteen *PEBP* genes (Liu *et al*., 2023a) in line with the observed expansion in other monocot lineages.

Seagrasses are foundation species and broadly are key components of marine coastal ecosystems essential for nutrient cycling, sediment stabilization, and habitats for fish and invertebrates (Orth *et al*., 2006; Hays *et al*., 2021). However, seagrasses are highly sensitive to natural and anthropogenic pressures such as eutrophication, shading, and direct bed disturbances, and experience higher mortality rates with higher water temperatures (Orth *et al*., 2006; Nejrup & Pedersen, 2008). Flowering and seed production play a key role in contributing to persistence and increasing genetic diversity within local *Z. marina* populations, and restoration efforts centered around seeds appear promising (Marion & Orth, 2010; Kendrick *et al*., 2012; Orth *et al*., 2012, 2020; van Katwijk *et al*., 2016; Cronau *et al*., 2023). Annual and perennial forms of eelgrass exist and are largely regarded as distinct ecotypes. In annual ecotypes, seeds germinate and shoots flower within one season (Robertson & Mann, 1984; Ruesink *et al*., 2022), whereas in perennial ecotypes, the predominant form, shoots persist for multiple seasons and populations as a whole produce far fewer flowering shoots with large variation in flowering frequency (Ruesink *et al*., 2022). Yet eelgrass shows large phenotypic variation, especially in flowering timing and frequency across spatial scales and seasons (Thom *et al*., 2003; Yang *et al*., 2013; Qin *et al*., 2020; Ruesink *et al*., 2022). Determining the causes for this variation is central to understanding the ecology, evolution, and restoration of the species.

In perennial populations of *Z. marina* in Willapa Bay, less than 25% of shoots typically flower each year (Thom *et al*., 2003; Ruesink *et al*., 2022), and it is not currently possible to predict how many and which shoots will flower. Therefore, studying the genetic and molecular mechanism of flowering onset in perennial populations proves very difficult. In the annual ecotype, all shoots flower in the same year as germination, such that populations dominated by annuals can exceed 70% flowering frequency (Phillips *et al*., 1983; Keddy, 1987; Ruesink *et al*., 2022). Characterizing the mechanism underpinning sexual reproduction and its relationship to environmental stimuli in *Z. marina* is a key knowledge gap that if addressed, would improve our understanding of how seed production is affected by local environmental factors. Further, identifying genetic markers of flowering would provide a method for predicting seed potential within a population. Such advances could ultimately allow for more strategic and efficient seed-based restoration efforts of eelgrass, through improved donor site selection and in the scaling-up of seed collection and planting efforts (Harwell & Rhode, 2007; van Katwijk *et al*., 2016; Orth *et al*., 2020). However, the relevant genes and mechanism by which *Z. marina* flowering onset is cued and regulated across populations remains completely unknown.

Here, we explored the mechanism of flowering in *Z. marina* and investigated the function of *FT/TFL1* homologs. We confirmed thirteen eelgrass florigen homologs (*ZmaPEBP*) and demonstrated that four are likely functionally relevant to the flowering pathway, both as activators and repressors. To elucidate *ZmaPEBP* gene function, we performed a heterologous functional assay in *Arabidopsis thaliana*. Overexpression of four *ZmaPEBP* genes, *ZmaFT2*, *ZmaFT4*, *ZmaFT9*, and *ZmaTFL1a,* resulted in either precocious or delayed flowering. We characterized expression patterns of these four genes in *Z. marina* shoots across three perennial sites and three annual sites to rule out patterns that are site- or life history-specific. Quantification of expression of these four genes across different tissue types and developmental stages suggests that *ZmaFT9* may play a role in the repression of flowering and *ZmaFT2* and *ZmaFT4* may contribute to flowering onset, while *ZmaTFL1a* likely functions in flowering shoot architecture.

## MATERIALS & METHODS

### Phylogenetic analysis and identifying *ZmaPEBP* genes

To identify *PEBP* family members and possible *FT* homologs in *Z. marina*, we performed a BLAST search (Altschul *et al*., 1990) using the *Arabidopsis* FT (AT1G65480) amino acid sequence against the *Z. marina* genome (Olsen *et al*., 2016). DNA sequences of *PEBP* family genes were obtained from the literature (Table S1), or from the genomes of the order *Alismatales* via a BLAST search with the *A. thaliana FT* and *Z. marina FT-like* sequence as queries. Sequences included previously described *PEBP* family genes from *A. thaliana*, *Oryza sativa, Aquilegia coerulea, Chrysanthemum seticuspe,* and *Brachypodium distrachyon* along with all recovered full-length *Alismatales* family *PEBP* genes, including *Z. marina*. To gain insight into the evolution and expansion of the *PEBP* gene family within other *Zostera* species, we included *Zostera muelleri* (Lee *et al*., 2016). *Z. muelleri PEBP* genes were identified via local BLAST search in its genome using *A. thaliana FT* sequences as query.

To create a phylogenetic tree, all full-length *Alismatales* sequences were aligned with previously described *PEBP* family sequences from *A. thaliana*, *O. sativa, A. coerulea, C. seticuspe,* and *B. distrachyon* by Multiple Sequence Comparison by Log-Expectation (MUSCLE) on the EMBL-EBI portal (Madeira *et al*., 2024). The alignment was fine-tuned by hand in Mesquite v 3.70 with the Multiple Sequence Alignment module (Maddison & Maddison, 2023). We used IQ-TREE (Nguyen *et al*., 2014) to build a maximum-likelihood phylogenetic tree with 1000 bootstraps using ModelFinder (Kalyaanamoorthy *et al*., 2017) to predict the best substitution matrix and UFBoot (Hoang *et al*., 2018) to compute ultra-fast bootstrap values. Mesquite Version 3.70 was used to build the consensus tree out of 400 base trees including sequences from *Z. marina, Z. muelleri, A. thaliana, B. distachyon, A. coerulea, Lemna aequinoctialis, Symplocarpus renifolius, Spirodela polyrhiza, C. seticuspe, O. sativa*, and *Allium cepa*.

Key amino acid residues that have been important in canonical FT and TFL1 structure and function (Hanzawa *et al*., 2005; Ho & Weigel, 2014) were analyzed using a multiple sequence alignment (MAFFT) (Katoh *et al*., 2002) of amino acid sequences of *Z. marina*, *Arabidopsis* FT and TFL1 along with the rice *Oryza sativa* FT homolog Hd3a to have a monocot FT sequence as comparison.

### Generating transgenic *ZmaPEBP* overexpression lines in *Arabidopsis* and quantifying flowering time

We used transgenic heterologous assays in *A. thaliana* to assess effect of *ZmaPEBP* genes on flowering and gain insights into overall function. All experiments in *A. thaliana* were performed in the wild-type Col-0 accession. To generate *35S:ZmaPEBP* transgenic lines, the coding sequences for each *ZmaPEBP* gene from the *Z. marina* reference genome (Olsen *et al*., 2016) were synthesized (Twist Bioscience, South San Francisco CA) and amplified with locus-specific primers by PCR to remove any adapter sequences (Table S2, S3). Using Gateway cloning, each gene was cloned into the pENTR D-TOPO vector (Invitrogen, Waltham MA), and after the insert sequences were verified by sequencing, they were transferred to the overexpression binary vector, pB7WG2 (Karimi *et al*., 2002). We used *Agrobacterium tumefaciens* strain GV3101 harboring each vector construct to transform *Arabidopsis* wild-type (Col-0) plants with conventional floral-infiltration and selection methods (Weigel & Glazebrook, 2002). The *35S:GFP* line was generated in a similar manner to serve as a control.

To screen for altered flowering phenotypes in *35S:ZmaPEBP* transgenic lines, we used T_1_ generation plants in the initial flowering time assay with all *35S:ZmaPEBP* and *35S:GFP* transgenic lines. In the T_1_ flowering time assay, sterilized seeds were germinated on selection media [1xLinsmaier-Skoog (PhytoTech Labs, Lenexa KS) media, 2% sucrose (wt/vol), 100 mg/L ticarcillin, 16 μg/mL Basta] in long day (LD) condition (16 h light, 8 h dark, 100 μmol/m^2^s, 22°C). 14-day-old seedlings with resistance to selection medium were transferred to soil (Sunshine #4, Sun Gro Horticulture, Agawam MA) and moved into plant growth chambers with LD conditions (16 h light, 8 h dark, 100 μmol/m^2^/s, 22 °C, 70% relative humidity). Flowering assays in the T_1_ generation included at least 16 individuals per construct. We recorded rosette leaf number at time of flowering.

Candidate genes that yield altered flowering time phenotypes were assayed using homozygous T_3_ plants with a single T-DNA insertion from *35S:ZmaPEBP* lines. In the T_3_ flowering assays, sterilized seeds were sown directly into soil in plant growth chambers with LD conditions (16 h light, 8 h dark, 100 μmol/m^2^/s, 22 °C, 70% relative humidity). We quantified flowering time in a similar manner in T_1_ and T_3_ lines. Flowering assays in the T_3_ generation included at least 10 individuals per line.

### Analyses of population genetic structure

In order to gain insight into the conserved nature of *ZmaPEBP* genes between populations and life history types, we described the genetic relatedness between meadows with annual and perennial ecotypes in close proximity to each other using sites in Willapa Bay, WA, USA. This well-studied estuary, located in the eastern Pacific Ocean, covers 240 km^2^ at mean tide (Hickey & Banas, 2003) and contains 34 km^2^ of eelgrass meadows occupying about a quarter of intertidal flats (Ruesink *et al*., 2006). The bay contains *Z. marina* populations comprising both the perennial and annual ecotype, which show widespread phenotypic and morphological variation, including temporal and population-specific variation in flowering frequency (Thom *et al*., 2003; Ruesink *et al*., 2022).

Population-level genetic structure was determined using reduced representation sequencing across five sites (RAD-seq; Figure 3 G, Table S4) (Ali *et al*., 2016). We chose three perennial and two annual sites, where one site included both ecotypes in close proximity (<0.2 km separation) (Table S4). In April 2023, 50 shoots were collected from Bay Center and Long Island, with 1-2 m spacing between each individual in order to reduce the probability of collecting clones (Duffy *et al*., 2022). The meristem region (3-5 cm) was saved from cleaned shoots and frozen in liquid nitrogen and stored at -80°C (Briones Ortiz *et al*., *In press*). Samples from Stackpole, Stackpole Annual, and Nahcotta Port Annual sites were collected in 2019 as described in Briones Ortiz et al (Briones Ortiz *et al*., *In press*).

High-molecular-weight genomic DNA was extracted using DNeasy Plant Pro Kit (Qiagen, Hilden, Germany). Genomic libraries were constructed for restriction-site associated DNA sequencing (RAD-seq) (Ali *et al*., 2016) with the restriction enzyme *Sbf1* and sequenced using the Illumina NovaSeq 6000 SP platform using 150 bp paired-end reads (University of Oregon, Genomics and Cell Characterization Core Facility, Eugene OR). Pooled-library reads were demultiplexed using the *process_radtags* program with *–best-rad* settings and trimmed to 137 bp in STACKS (Catchen *et al*., 2013). The single-end reads were aligned to the published genome of *Z. marina* (Olsen *et al*., 2016; Ma *et al*., 2021), using BWA-MEM (Li, 2013), with minimum alignment and mapping Phred quality scores of 30.

Genotyping of polymorphic loci was conducted using the *ref_map* pipeline in STACKS (Rochette & Catchen, 2017). Subsequent filtering was performed using the R-package SNPFILTR (DeRaad, 2022)(steps described in Table S5). Identical multilocus genotypes (MLGs, >95% shared alleles between individuals), a result of clonal reproduction, were identified using PLINK (function –genome) (Chang *et al*., 2015). Only one representative sample of each clonal genotype was retained for further processing. Samples in the Stackpole perennial and both annual sites were initially collected as seedlings resulting from sexual reproduction, and no clones were found.

Population structure was examined using measures of population genetic differentiation (F_ST_) (Weir & Cockerham, 1984) using the R-package HIERFSTAT (Goudet, 2005; de Meeûs & Goudet, 2007). Population structure was also inferred using a test of population assignment of individuals, implemented in STRUCTURE v2.1.1.5 (Pritchard *et al*., 2000). The number of clusters (*K*) tested ranged from 1 to 10, and the most likely ancestral groups were evaluated using the mean log probability of the data, *L*(*K*) (Pritchard *et al*., 2000) and the *ΔK* statistic (Evanno *et al*., 2005). Genetic relationships among individual samples were visualized using a discriminant analysis of principal components (DAPC) plot, implemented in the R package ADEGENET (Jombart & Ahmed, 2011).

### Collection of plant material, RNA extraction and quantitative RT-PCR

We used extracted RNA and quantitative RT-PCR (qPCR) to assess gene expression levels in both *Z. marina* and *A. thaliana* plant tissue. Gene expression was determined in three instances: *A. thaliana* T_1_ and T_3_ lines, and *Z. marina* in field collections. For gene expression analysis in T_1_ lines, sterilized seeds were germinated on 1xLS 2% sucrose antibiotic selection media in LD conditions (16 h light, 8 h dark, 100 μmol/m^2^s, 22 °C) to isolate successful transformed seedlings. At two weeks, seedlings were transferred to 1xLS media without sucrose and antibiotics. After one week, seedlings (21-day-old) were harvested at ZT16 (zeitgeber time). For gene expression analysis in T_3_ lines, seeds were germinated on 1xLS media without sucrose in LD conditions (16 h light, 8 h dark, 100 μmol/m^2^s, 22 °C). 14-day-old seedlings were collected at ZT16.

For gene expression analysis in *Z. marina* from the field, whole shoots and plant tissues (including rhizome and roots, vegetative mature leaves, inflorescence, and spathes) were collected from perennial population sites in Willapa Bay (Fig. 3, Table S4) in a three-consecutive-day period in mid-July (13-15 Jul 2022) between ZT3-5. We collected tissue from at least 15 adult plants at each site, with 1-2m spacing between each individual. Samples from two annual populations [ST-ANN and NP-ANN, (Ruesink *et al*., 2022)] and one annual population from Yaquina Bay (YQ-ANN) were collected in a similar manner at two-week intervals between April and August 2023 between ZT3-5. All plant tissue collected was stored immediately in RNAlater stabilization solution (ThermoFisher, Waltham MA). Samples were moved to -80°C for long-term storage. Photoperiod data was obtained from the US Naval Astronomical Applications Department (Astronomical Applications Department, U. S. Naval Observatory, 2023). Temperature data was collected at the sediment surface using iButtons (Dallas Semiconductor, Dallas TX) in 2-hour increments at Willapa Bay Annual sites.

Total RNA was isolated from samples using RNeasy Plant Mini Kit (Qiagen, Hilden, Germany), including an on-column DNA digestion procedure. Complementary DNA (cDNA) was obtained with iScript cDNA synthesis kit (Bio-Rad, Hercules CA). We analyzed expression levels of the gene of interest with qPCR. The qPCR reaction was carried out on 2μL of 1/5-diluted cDNA using 2X SSoAdvance SYBR Super Mix (Bio-Rad, Hercules CA) with locus-specific primers (Table S2).

We used two reference genes to normalize expression in transgenic *Arabidopsis*, *ISOPENTENYL PYROPHOSPHATE / DIMETHYLALYLL PYROPHOSPHATE ISOMOERASE* (*IPP2*) and *SERINE/THREONINE PROTEIN PHOSPHOTASE 2A* (*PP2A*) (Song *et al*., 2018). We used three previously described reference genes for expression analysis in *Z. marina* (Ransbotyn & Reusch, 2006), *CYCLOPHILIN 2* (*CYP2*), *EUKARYOTIC INITIATION FACTOR4A* (*ELF4A*), and *RIBOSOME STRUCTURAL PROTEIN L28* (*RPL28*). Samples with no detectable expression were treated as having a cycle threshold (C_T_) value as C_T_=40. A baseline threshold of relative fluorescence units (RFU) was set for consistency across primers to compare between plates, and resulting C_T_ values were analyzed using a delta C_T_ method. For the reference, the average C_T_ values of all reference genes were used for calculation.

### Statistical analysis of *Z. marina* gene expression data

We performed statistical analyses on the *Z. marina* expression data to compare tissue-specific expression in flowering and vegetative life stages across three sites (Jul 2022), and to examine the time series of expression in annual eelgrass (Apr-Sep 2023). Expression of candidate genes (*ZmFT2*, *ZmFT4*, *ZmFT9*, and *ZmTFL1a*) were analyzed separately. In the first comparison, linear models (analysis of variance, ANOVA) were constructed with relative expression values (delta C_T_) as response variable and with site, tissue, and life stage, as well as tissue x life stage interaction included as fixed factors. To be comparable between vegetative and flowering life stages, the included tissue types were rhizome, stem, and leaf middle (middle 10cm of leaf blade, Fig. 3).

In the second comparison, each of the eight collection timepoints was considered a separate level of a categorical factor, because different individuals were collected each time, and because gene expression could turn on or off abruptly. Using a linear mixed effects model framework, delta C_T_ was the response variable and fixed effects were time point and site. Additionally, we tested for a site by time point interaction to that would account for different phenology between sites. Expression values required log-transformation to generate a Gaussian distribution in all analyses. Any significant factors or interactions were followed up by post-hoc tests to determine significant differences between groups based on Tukey HSD significance levels. Statistical models were built in R (R Core Team, 2024) using the package lme4 (Bates *et al*., 2015), with post-hoc tests from emmeans (Searle *et al*., 1980).

## RESULTS

### Phylogenetic analysis revealed thirteen *PEBP* genes in *Z. marina*

We identified thirteen *Z. marina* genes that fell within the *PEBP* gene family based on sequence homology. Phylogenetic analysis incorporating other *PEBP* genes representing different plant lineages revealed these thirteen genes belong to three clades; *FT/TSF* (ten genes, named *ZmaFT1-10*), *TFL1/BFT* (two genes, labeled *ZmaTFL1a* and *ZmaTFL1b*), and *MFT* (one gene, *ZmaMFT1*) (Fig. 1). Monocot lineages harbor an expansion in *FT* genes, and it has been shown that the initial *FT* duplication within monocots occurred in *Alismatales*, the basal monocot lineage that includes *Z. marina* and the *Zosteraceae* family (Bennett & Dixon, 2021). *Z. muelleri*, a sister species to *Z. marina*, has a similar gene duplication structure to *Z. marina*. Both *Z. marina* and *Z. muelleri* have multiple copies of *FT* beyond the initial duplication described in Bennett & Dixon (Bennett & Dixon, 2021) and have copies present in three *FT* subclades described in Liu et al. (Liu *et al*., 2023a).

**Figure 1:**
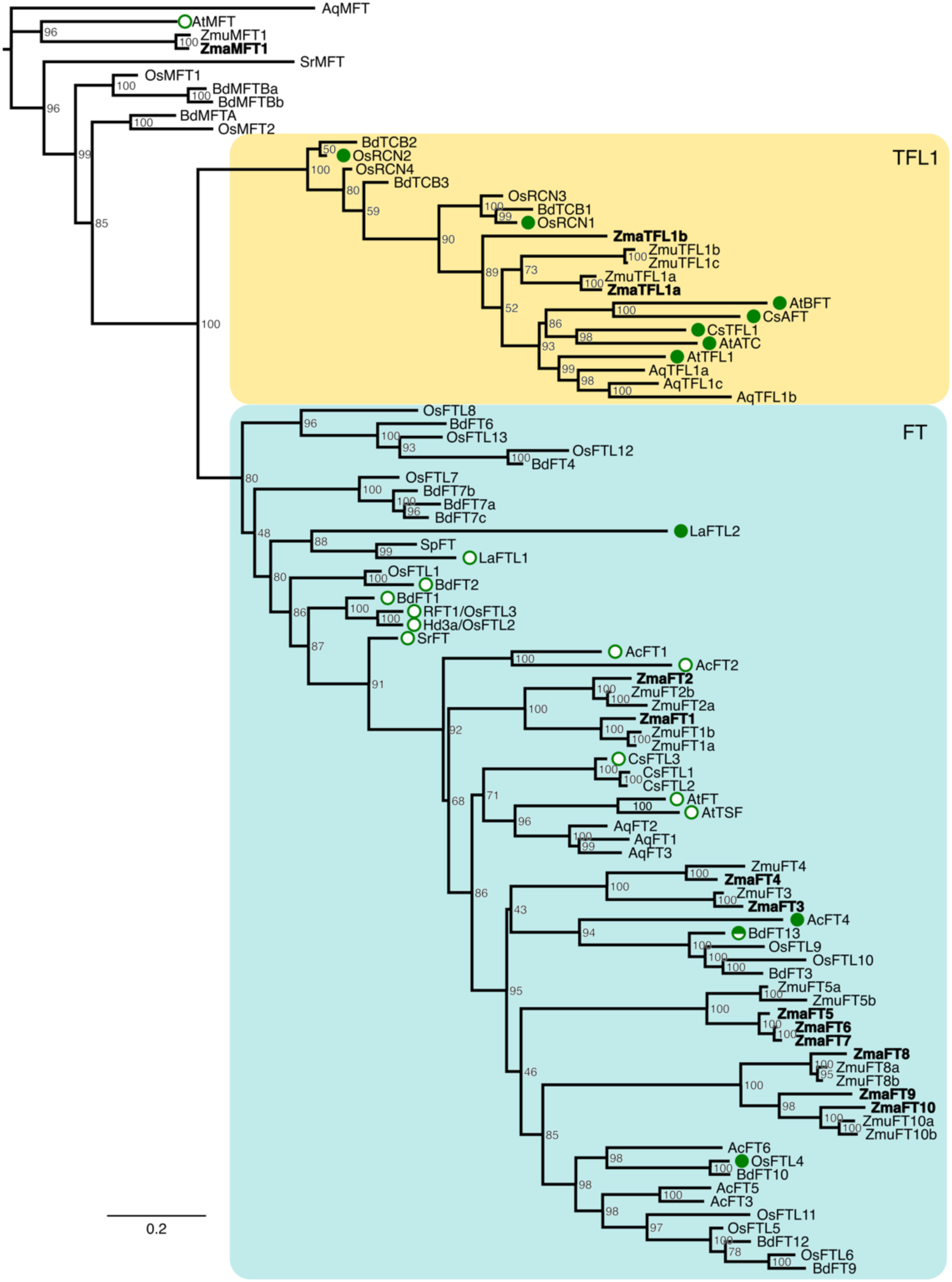
Phylogenetic tree of phosphatidylethanolamine binding protein (PEBP) genes in angiosperms. See Table S1 for sequence accession numbers. Nucleotide-level maximum likelihood analysis with 100 bootstraps, values shown at nodes. Species were chosen to represent the different angiosperm lineages having *PEBP* gene family members, with over-representation of monocotyledoneous and *Alismatales* taxa (monocotyledons: *B. distachyon, O. sativa*, *A. cepa*; *Alismatales*: *L. aequinoctialis, S. polyrhiza, S. renifolius, Z. marina, Z. muelleri;* other: *A. thaliana, A. coerulea, C. seticuspe*). *Z. marina* characterized here is shown in bold, and the tree includes its sister species *Z. muelleri* (*Zmu*). *PEBP* genes fall into two clades: *FT* (*FLOWERING LOCUS T,* blue) and *TFL1* (*TERMINAL FLOWER 1*, yellow). Open circles denote previously described activators of flowering, closed circles denote repressors, and half-filled circle denotes activator (under short-day conditions) and repressor (under long-day conditions).

*PEBP* genes that activate flowering tend to cluster within the *FT* clade, while genes that repress flowering tend to cluster with the *TFL1* clade (Pin *et al*., 2010; Bennett & Dixon, 2021; Liu *et al*., 2023a). Our analysis showed *Z. marina* and *Z. muelleri* both have genes that cluster within the *FT* clade and *TFL1* clade, as well as one predicted *MFT* paralog each. While sequence similarity cannot confirm or predict gene function (Delaux *et al*., 2019), our analysis suggest that the *Z. marina* genome may have several representatives of the same floral pathway genes that are conserved across angiosperms.

### Four *ZmaPEBP* genes affect flowering phenotype in overexpression assays

As a first step towards investigating the function of the *ZmaPEBP* genes, we characterized their protein structure and function. The predicted open reading frames for all *ZmaPEBP* genes encode proteins with high similarity to *Arabidopsis* FT and TFL1 of *Arabidopsis* and rice Hd3a of *Oryza sativa* (Fig. 2a, Fig. S1). We also observe the conservation of key residues for FT function (Hanzawa *et al*., 2005; Ho & Weigel, 2014), such as at position 85 where all genes show conservation relative to *Arabidopsis* FT and TFL (Y85 in FT, and H85 in TFL1).

**Figure 2:**
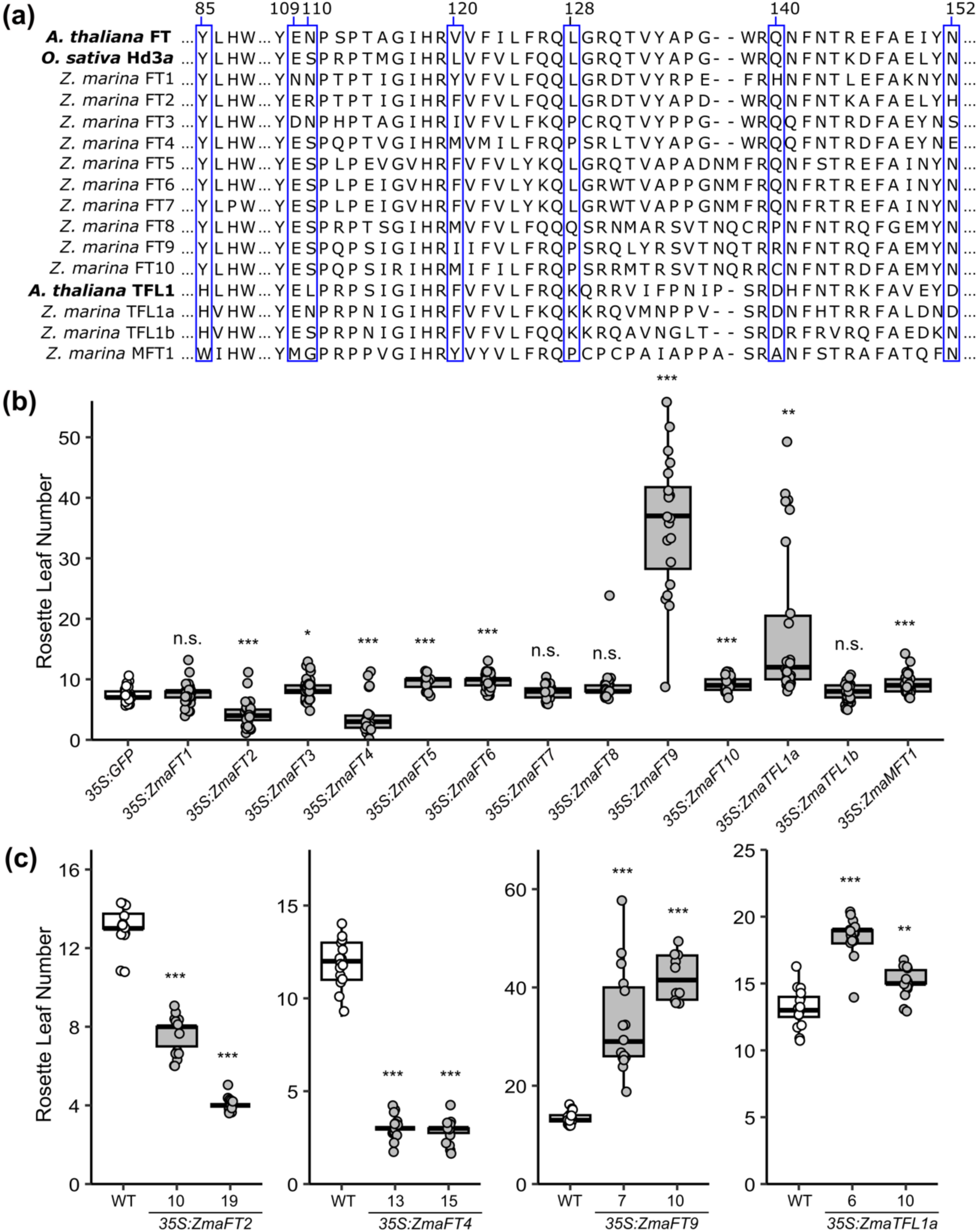
*ZmaFT* and *ZmaTFL1* genes alter flowering phenotype when overexpressed in *Arabidopsis thaliana.* **(a)** Amino acid sequence alignment of ZmaFT and ZmaTFL1 with *Arabidopsis thaliana* FT and TFL1 and *Oryza sativa* Hd3a (an FT ortholog). Blue boxes highlight residues important for FT function as a flowering promoter (blue). **(b)** Overexpression of thirteen *ZmaPEBP* orthologs under the control of the *35S* promoter highlighted four candidates that altered flowering time. *ZmaFT2* and *ZmaFT4* cause precocious flowering, while *ZmaFT9* and *ZmaTFL1A* caused delayed flowering in the T_1_ generation under long-day conditions. Asterisks indicate significance based on t-test against the control (*35:GFP*), with Bonferroni correction (n.s.: non-significant, *: *p* < 0.0038, **: *p* < 0.00077, ***: *p* < 0.000077). **(c)** Altered flowering time phenotype was confirmed in homozygous individuals (T_3_ generation). Number labels on x-axis represent homozygous lines. Asterisks indicate significance based on t-test comparing to wild type (WT) with Bonferroni correction (**: *p* < 0.005, ***: *p* < 0.0005). All individuals are shown as data points (n≥16), with median indicated by center line in box, upper and lower quartile by box boundaries, and highest and lowest value within two interquartile ranges by whiskers.

To gain further insight into the function of each *ZmaPEBP* gene, we created transgenic overexpression *Arabidopsis* lines (Col-0) of each *ZmaPEBP* gene to investigate the effect of expression of each candidate gene on flowering *Arabidopsis*. We validated *ZmaPEBP* expression in the T_1_ generation (Fig. S2). Of the thirteen *ZmaPEBP* genes screened, four demonstrated a strong effect on flowering time. Overexpression of *ZmaFT2* and *ZmaFT4* resulted in early flowering in *Arabidopsis* under LD conditions, compared to a *35S:GFP* control (Fig. 2b). Both genes are found in the *FT/TSF* clade in our phylogenetic analysis (Fig. 1). In homozygous transgenic lines, overexpression of *ZmaFT4* had a stronger effect on flowering time than *ZmaFT2* (Fig. 2c, Figure S3a). *ZmaFT9* and *ZmaTFL1a* lines showed delay in flowering time, with a much stronger phenotype observed in *ZmaFT9* lines in homozygous lines (Fig. 2b-c). *ZmaTFL1a* is in the *TFL/BFT* clade with *Arabidopsis TFL1*, while *ZmaFT9*, which has the stronger delay in flowering time, is found in the *FT/TSF* clade (Fig. 1). We also observed a decrease in *LFY* and *API*, downstream targets of *FT*, in homozygous *35S:ZmaFT9* and *35S:ZmaTFL1a* lines, indicating some effect on the *FT* pathway in *Arabidopsis* (Fig. S3b). Further, flowers produced on T_1_ generation plants showed a morphological phenotype similar to *TFL* overexpression *Arabidopsis* plants and to mutant *lfy* plants where additional leaf-like structures develop in place of petals and floral organs (Huala & Sussex, 1992; Hanano & Goto, 2011) (Fig. S3c). Together, these results indicate that *Z. marina* has two *PEBP* genes that promote flowering and two that repress flowering, all of which do so through interaction with the floral pathway.

### Genetic structure between annual and perennial ecotypes provides context for the potential roles of *ZmaPEBP* genes

To study the roles of *ZmaFT2*, *ZmaFT4*, *ZmaFT9*, and *ZmaTFL1a* in eelgrass flowering and shoot architecture in plants with varying genetic backgrounds, we analyzed gene expression in vegetative and flowering shoots across different natural populations. When eelgrass flowers, the vegetative shoot primarily made up of leaves originating at the shoot base produces a bolted stem with multiple inflorescences, completely changing the shoot architecture of the plant (Fig. 3a-d). We focused on sampling plants from eelgrass meadows with shoots undergoing both clonal and sexual reproduction at three perennial sites, and from meadows where sexual flowering is the predominant mode of reproduction at three annual sites (two within Willapa Bay and one in Yaquina Bay; Fig. 3e-g, Table S4).

**Figure 3:**
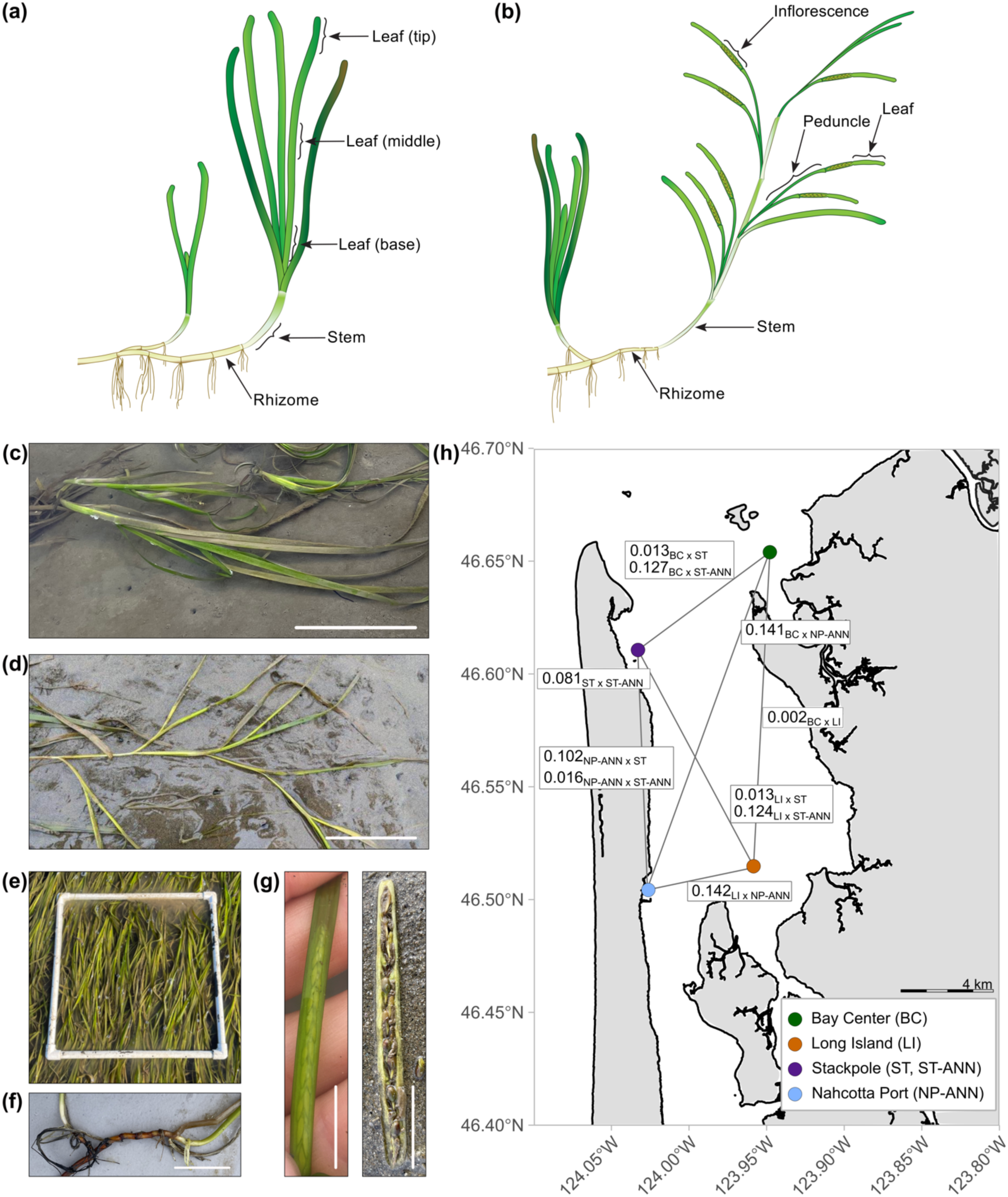
Description of *Zostera marina* (eelgrass) sampling approaches, life stages, and sites used in this study. Tissue isolated from **(a)** vegetative and **(b)** flowering shoots. Eelgrass **(c)** vegetative and **(d)** flowering shoots. Scale bar approximately 10 cm. **(e)** Eelgrass meadows form through **(f)** clonal reproduction via rhizome extension or **(g)** from the formation of inflorescences and seeds within spathes. Scale bar approximately 1 cm. **(h)** Geographic location of study sites in Willapa Bay, WA USA: Bay Center (BC), Long Island (LI), Stackpole (ST), Stackpole Annual (ST-ANN) (ST and ST-ANN represented by sample point on map due to proximity), and Nahcotta Port Annual (NP-ANN) (Table S4). Pairwise genetic distance, F_ST_, estimated through estimated RAD-Seq analysis shown in white boxes (n=48 per population). Subscript of F_ST_ indicates which sites included in the pair-wise comparison. Scale bar approximately 4 km. Photographs of annual plants are in Supplemental Fig. S6.

To test whether population structure or life history type influenced expression patterns in *ZmaPEBP*, we estimated genetic distance and population structure across these five eelgrass meadows (Fig. 3e-g). Reduced representation sequencing (RAD-seq) identified 327 loci (single nucleotide polymorphisms; SNPs) across 224 individuals (Table S4). Tests for population structure (Fig. 3h, Fig. S4, Table S6) revealed significant genetic differentiation between life history types (annual and perennial), and, to a lesser extent, between geographic regions within ecotypes. Pairwise genetic distances (F_ST;_ Table S6) were highest and significant for all pairwise comparisons between annual and perennial sites, including the co-located annual and perennial sites at Stackpole. Amongst the perennial populations, the Stackpole perennial population in the west of the Bay was significantly differentiated from the two populations in the east, but these distances were lower than between annual and perennial populations; however, there was no significant structure between the eastern populations, Bay Center and Long Island (Table S4). Small but significant genetic differentiation was also observed between the annual sites, Stackpole annual in the north and Nahcotta Port in the south (Table S4). Similarly, DAPC analyses (Fig. S4b) separated annual from perennial populations on the first axis (explaining 71.3% of the variation in the data set) and both north-south (annual sites) and east-west (perennial sites) geographic structure on the second axis (12.3% of the variation). Finally, tests for individual population assignment revealed two primary clusters (*K* = 2), explained primarily by the assignment of most individuals in the annual population one group, and those from perennial populations to the second group (Fig. S4A). Overall, population structure across ecotypes, as well as across small geographic distances within ecotypes, (less than 20km in a single bay) allowed us to test for common gene expression patterns associated with flowering. Finally, we also included annual samples from Yaquina Bay (Table S4), 140 km to the south and likely genetically distinct (Yu *et al*., 2023) in the expression analyses.

### Expression of *ZmaPEBP* genes changes across development and tissue type

To gain insight into the roles of *ZmaFT2*, *ZmaFT4*, *ZmaFT9*, and *ZmaTFL1a* in eelgrass flowering, shoot architecture and development, we analyzed gene expression in different tissues from both adult vegetative and flowering shoots from each perennial population (Fig. 4a-b, Table S4). In *Z. marina*, the two flowering activators, *ZmaFT2* and *ZmaFT4* (Fig. 2b-c), had higher expression in stem and rhizome of flowering compared to vegetative shoots, but expression in leaves was more similar between life stages (Fig. 4b, Fig. S5, Table S7). Although *ZmaTFL1a* acted as a flowering repressor when overexpressed in *Arabidopsis* (Fig. 2b-c), its expression decreased in flowering tissue relative to vegetative shoots, whereas expression in stem and rhizome tissue increased in flowering relative to vegetative shoots (Fig. 4a-b). This result was surprising, since the shoot apical meristem of the plant was in the vegetative stem tissue section, and expression was observed in tissues beyond the shoot apical meristem, where *TFL1* expression is typically found in other plant species (Bradley *et al*., 1997; Ratcliffe *et al*., 1999; Nakagawa *et al*., 2002; MacAlister *et al*., 2012). Both *ZmaFT4* and *ZmaTFL1a* showed significantly higher levels of expression in the tip of vegetative leaves compared to other sections of vegetative leaves (Fig. S5, Table S8). These spatial distribution patterns within the leaves resembled that of *FT* in *Arabidopsis* (Takada & Goto, 2003). The expression levels of the other floral repressor, *ZmaFT9* (Fig. 2b-c), were lower in leaves and stems from flowering shoots compared to vegetative shoots (Fig. 4a-b, Fig. S5, Table S3). Together, these results suggest that *ZmaFT9* correlates to the vegetative state in eelgrass adult perennials.

**Figure 4:**
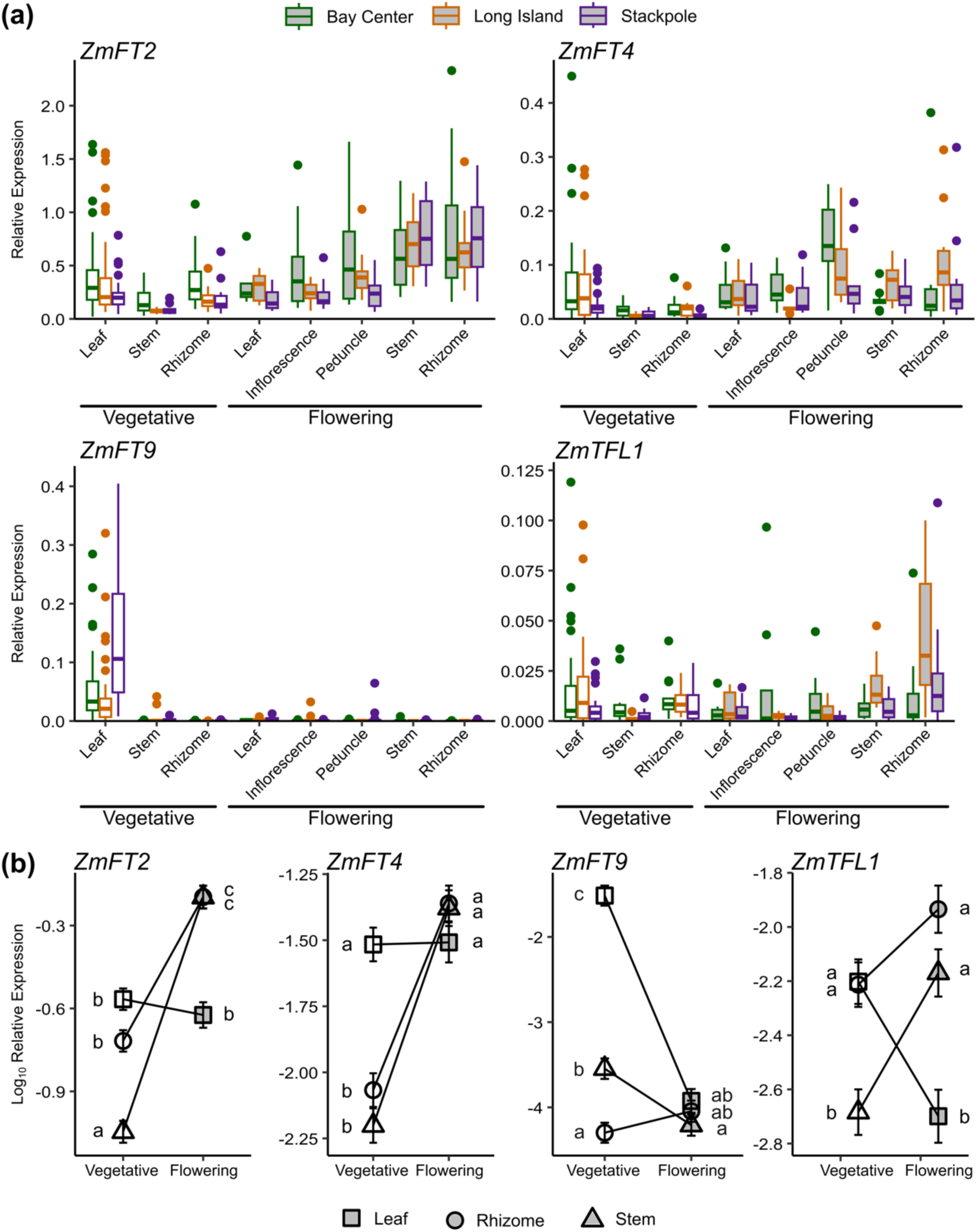
Expression of florigen genes in different *Z. marina* tissues. **(a)** Relative expression levels of *ZmaFT2*, *ZmaFT4*, *ZmaFT9*, and *ZmaTFL1a* across different tissue types in different developmental stages, vegetative (white) and flowering (grey). Expression in tissues was measured in samples (n≥10) from 3 sites shown in Figure 3H. All expression values are relative to 3 reference genes (*CYP2*, *ELF4A*, and *RPL28*). Median is indicated by center line in box, upper and lower quartiles by box boundaries, and highest and lowest values within two interquartile ranges by whiskers. **(b)** Comparing means between tissue types across reveals different trends of expression changes within tissue types across vegetative (white) and flowering (grey) developmental stages. Mean expression was calculated using log_10_ transformed expression values. Error bars are standard errors of means. Different letters indicate statistically significantly different groups determined by one-way ANOVA test with post-hoc Tukey HSD of log_10_ transformed values.

Our results in *Z. marina* highlight the expression of floral activators in stem and rhizome tissue, a pattern not observed in *Arabidopsis*. *ZmaTFL1a*, contrary to its apparent function as a floral repressor in *Arabidopsis*, is upregulated in stem and rhizome flowering tissue. However, *TFL1* is known to also play a role in shoot architecture and development in *Arabidopsis* (Shannon & Meeks-Wagner, 1991; Kobayashi *et al*., 1999), which aligns with our observed expression in flowering stems. Overall, our results suggest that *ZmaFT2*, *ZmaFT4*, *ZmaFT9*, and *ZmaTFL1a* are implicated in eelgrass floral development and show tissue-specific expression patterns.

### Expression of *ZmaPEBP* genes changes over the lifecycle of an annual ecotype

Due to the annual ecotype’s predictability of flowering, annuals provide a system in which to study *PEBP* gene expression trends over the development of the plant throughout the growing season and their relationship to flowering onset. To this end, we characterized expression of *ZmaFT2*, *ZmaFT4*, *ZmaFT9*, and *ZmaTFL1a* at different time points across the growing season from two annual populations in Willapa Bay (Stackpole Annual and Nahcotta Port Annual, Fig. 3h). In the 2023 season in Willapa Bay, seedlings at these sites emerged in early April, and differentiation to flowering occurred in late June (Fig. 5a, Fig. S6). The emergence of flowering shoots coincides with peak photoperiod (approximately 16 hours, Fig. 5b).

**Figure 5:**
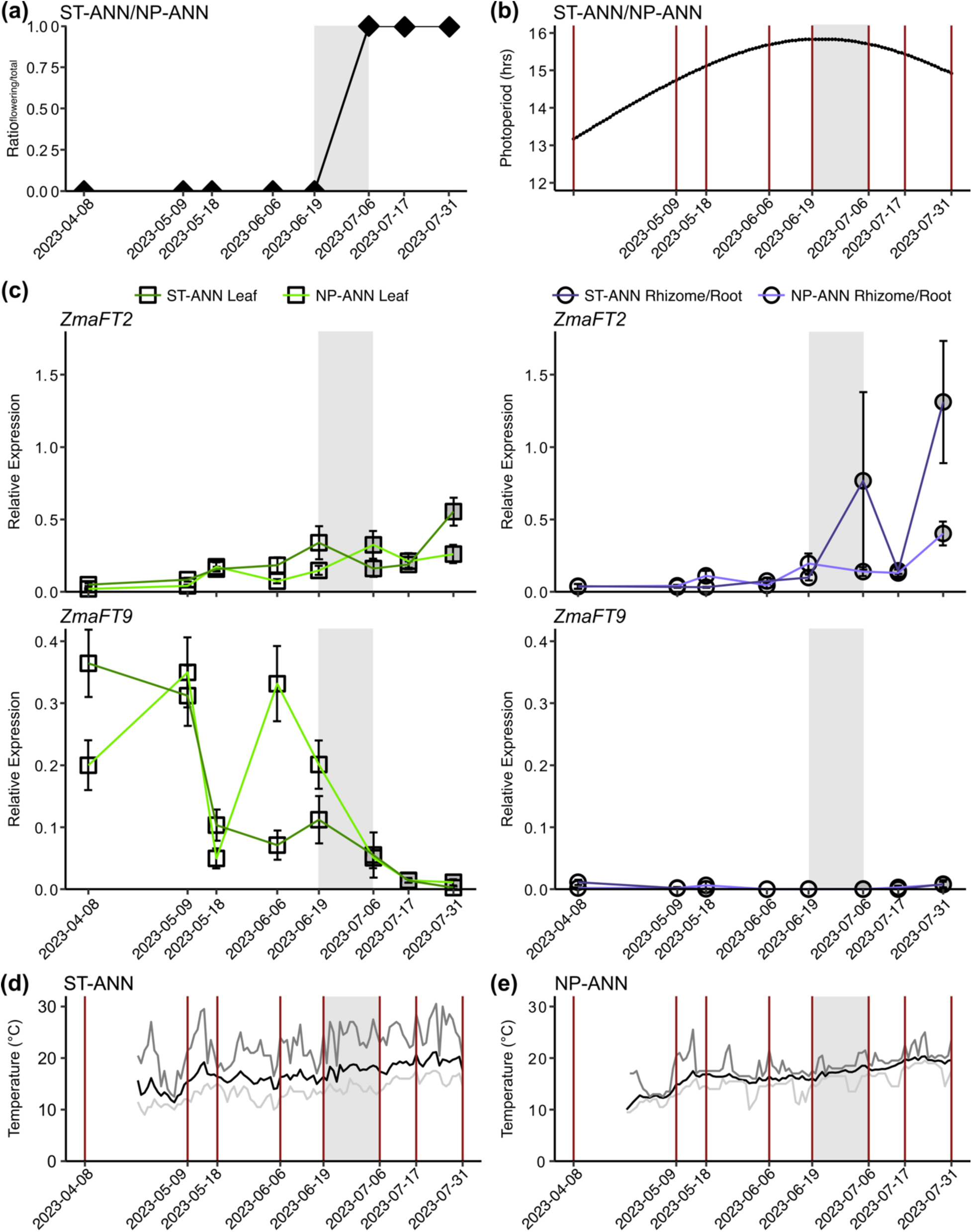
Expression of *ZmaFT2* and *ZmaFT9* genes in leaf and rhizome tissue across the 2023 growing season at ST-ANN and NP-ANN sites (Figure 3H, Table S4). **(a)** Ratio of flowering shoots to total shoots collected at each time point for both ST-ANN populations and NP-ANN population, n≥10. Grey shaded region in each panel indicates when flowering shoots emerged. **(b)** Photoperiod regime over growing season in Willapa Bay. Data obtained from the US Naval Astronomical Applications Department. Red lines indicate days samples were collected for gene expression analysis. **(c)** Relative expression levels of *ZmaFT2* and *ZmaFT9* across leaf (square) and rhizome and root (circle) throughout the growing season (April-July). Lighter green and purple lines indicate NP-ANN site, while darker green and purple lines indicate ST-ANN site. Time points where shoots are vegetative shown in white and time points where shoots had flowered are filled grey. All expression values are relative to 3 reference genes (*CYP2*, *ELF4A*, and *RPL28*). Plot point represents mean, and error bars are standard error. Statistically significant groupings based on post-hoc Tukey HSD of log_10_ transformed values are listed in Table S10. **(d, e)** Daily average temperature (black), daily minimum temperature (light grey), and daily maximum temperature (dark grey) at ST-ANN site **(d)** and NP-ANN site **(e)** throughout the growing season. Temperatures were collected at the sediment surface using iButtons (Dallas Semiconductor) every 2 hours. Red lines indicate days samples were collected for gene expression analysis. Temperature from 2023-04-08 to 2023-04-23 was not recorded.

In both *ZmaFT2* and *ZmaFT4*, we observed increasing expression over the season, with a more significant increase occurring when flowering shoots emerged (Fig. 5c, Fig. S7, Table S9, Table S10). This trend was far more apparent in root and rhizome tissue than in leaf tissue, though there was a marked increase in leaf tissue gene-expression on 19 Jun 2023 (Julian Date 23170) in ST-ANN and 6 Jul 2023 (Julian Date 23187) in NP-ANN, just as bolted flowering shoots were observed, which dissipated by the next time point. A similar trend was observed in *ZmaTFL1a* expression, with slight increased expression in leaves just as flowering shoots emerged and a general increase in expression after flowering shoots emerged, primarily in roots and rhizomes (Fig. S7). These results align with expression trends observed in perennial shoots (Fig. 4b). The peak in *ZmaFT2*, *ZmaFT4*, and *ZmaTFL1a* leaf expression also coincides with the maximum photoperiod (Fig. 5b). *ZmaFT9*, unlike the other *ZmaPEBP* genes, is expressed at higher levels in leaves at earlier stages of vegetative development. Approximately one month before flowering shoots were observed, we saw a decrease in expression (Fig. 5c). The average daily temperature rose 3°C (12°C to 15°C) and 1.9°C (14.9°C to 16.8°C) between 9 May 2023 (23129) and 18 May 2023 (23138) during this interval at ST-ANN and NP-ANN, respectively, with daily maxima ranging from 18°C to 29.5°C at ST-ANN and 17°C to 25.5°C at NP-ANN (Fig. 5d-e). The large decline in *ZmaFT9* expression was followed by a slower decline in expression after flowering shoots emerged. These results also align with tissue and developmental state-specific expression trends observed in perennial shoots (Fig. 4b). We also analyzed expression levels of *ZmFT2*, *ZmFT4*, *ZmFT9*, and *ZmTFL1a* in annuals from Yaquina Bay (YQ-ANN, Table S4). Trends in expression of each gene align with expression patterns observed in ST-ANN and NP-ANN site (Fig. S8) however results from YQ-ANN were either not statistically significant or less significant, likely due to limited sample size size and sampling of clonal branches rather than bolting portions (Table S11 and S12).

Together, these results suggest that *ZmaFT2* and *ZmaFT4* are involved in the activation of flowering, potentially in the formation of floral meristems and inflorescences, with *ZmaTFL1a* involved in some shoot architecture function after the flowering shoot has bolted. *ZmaFT9*, on the other hand, is seemingly involved with vegetative growth and development, and appears to require a decrease in expression to allow flowering onset to occur.

## DISCUSSION

In this study, we aimed to characterize florigen genes in eelgrass and to examine their function in regulating the onset of flowering. The molecular controls and regulatory mechanisms for the onset of flowering in *Z. marina* have not been previously investigated, despite the importance of sexual reproduction in improving resiliency in populations, a key goal for restoration efforts (Kendrick *et al*., 2012). Identifying and characterizing flowering genes and their effect on flowering onset in eelgrass will inform how flowering may be cued in eelgrass and how eelgrass is predicted to sexually reproduce under climate change conditions.

Here, we provide evidence of functional *FT/TFL1* homologs in *Z. marina*, with both activating and repressing function in flowering. We identify thirteen florigen homologous genes in eelgrass and demonstrate that four are functionally relevant to the flowering pathway. We show that *Z. marina*, like other monocots, has an expanded *FT* clade within the *PEBP* gene family, which is also mirrored in *Z. marina*’s sister species, *Z. muelleri*. Using *Arabidopsis* to assess the function of *Z. marina PEPB* genes, we demonstrate that overexpression of four *ZmaPEBP* genes results in either precocious flowering (*ZmaFT2* and *ZmaFT4*) or delayed flowering (*ZmaFT9* and *ZmTFL1a*). To the best of our knowledge, these results provide the first functional implication of florigen in marine angiosperms. Further, we observed tissue-specific expression patterns of these *ZmaPEBP* genes that correlate to developmental state (flowering and vegetative) in both annual and perennial *Z. marina* shoots. We believe that the role of the four focal genes (*ZmaFT2*, *ZmaFT4*, *ZmaFT9*, and *ZmaTFL1a*) is conserved across annual and perennial ecotypes, and across geographically structured populations, based on the similarity of gene expression patterns (Fig. 4 and 5). Our findings add to the expanding body of knowledge on seasonal flowering through the balance of floral activators and repressors and bring new insight into *FT*-flowering pathway function beyond terrestrial plants and model and crop species.

### *Z. marina* harbors a large expansion of *FT* genes with *ZmaFT2* and *ZmaFT4* likely contributing to flowering onset

In *Arabidopsis*, there are six *PEBP* genes (*FT*, *TSF*, *TFL1*, *ATC*, *BFT*, and *MFT*), with *FT* as the main inducer of flowering and *TSF* with redundant functionality. Our phylogenetic analysis demonstrated an expansion in the *PEBP* gene family focused within the *FT* clade. Ten of the thirteen *ZmaPEBP* genes identified fell within *FT/TSF* clade, consistent with the phylogenetic analysis of *FT* genes described in Liu et al. (Liu *et al*., 2023a). Further completion and annotation of the *Z. marina* genome may reveal other florigen genes as well as insights into other genes implicated in flowering processes. In eelgrass, *ZmaFT2* and *ZmaFT4* likely contribute to the activation of flowering in a manner similar to *FT* and are apparent drivers of flowering. Interestingly, *ZmaFT1*, the closest related paralog to *ZmaFT2*, shows no flowering phenotype when overexpressed in *Arabidopsis* (Fig. 2b). We speculate that this lack of apparent flowering function in *Arabidopsis* may be due to a mutation in ZmaFT1 at a key amino acid residue, Q140 (Ho & Weigel, 2014), which is a histidine (H) in ZmaFT1 rather than a glutamine (Q) but is conserved in both ZmaFT2 and ZmaFT4 (Fig. 2a). Gln140 forms hydrogen bonds with Tyr85 (Y85), another important residue (Hanzawa *et al*., 2005; Ho & Weigel, 2014), conserved in all ZmaFT homologs within the FT clade. In wild populations, both *ZmaFT2* and *ZmaFT4* show increased expression in stem and rhizome tissue after the onset of flowering in both perennial and annual shoots. Further, there is a slight increase in expression observed in leaf tissue just before the emergence of flowering shoots. This peak in expression in annual populations approximately coincided with maximum photoperiod, indicating that photoperiod may be one influencing factor on the expression of these *ZmaFT* genes. This result coincides with previous literature, where temperature, salinity, and photoperiod were found as environmental controls of flowering (McMillan, 1976; Harwell & Rhode, 2007; Blok *et al*., 2018).

### *ZmaFT9* may be the main determinant of flowering through repression of flowering function

Despite the importance of activators within the *PEBP* gene family and other plant species, our results suggest that a repressor, namely *ZmaFT9*, may play a role as a major determinant of flowering onset in *Z. marina*. Not only did overexpression of *ZmaFT9* have a significant delay in flowering time in *Arabidopsis*, but its expression was also restricted to vegetative leaves in both the annual and perennial populations in *Z. marina*. Moreover, *ZmaFT9* expression decreased sharply approximately one month before flowering shoots emerge in the annual ecotype. Only after *ZmaFT9* expression decreases do we observe increased expression in *ZmaFT2* and *ZmaFT4*, suggesting that they are likely activators of flowering. These results suggest the involvement of *ZmaFT9* in the repression of flowering and maintenance of the vegetative state. Given that *ZmaFT9* is found only in leaves, we hypothesize that *ZmaFT9* acts as an anti-florigen (Higuchi, 2018), which too is expressed in the leaves like canonical *FT* (Takada & Goto, 2003) but prevents activation of flowering through repression of flower-activating gene expression, and ultimately is a major determinant of flowering onset on *Z. marina*. In this framework, the presence of antiflorigen maintains the vegetative state in shoots, and when it is substantially reduced, the shoot experiences the onset of flowering. Adult perennial shoots that remained vegetative during peak flowering time in the field express *ZmaFT9* at high levels, which was not seen in adult perennial flowering shoots (Fig. 4). High expression levels of *ZmaFT9* throughout the time of flowering onset may explain the low flowering frequency observed in perennial populations.

Most *PEBP* genes within the *FT* clade activate flowering in a similar manner to the florigen described originally in the 20^th^ century, where a floral stimulus originates in the leaves and travels to the shoot meristem (Chailakhyan, 1937). However, a *FT* homolog that functions antagonistically to a canonical florigen is not completely novel. In onions (*Allium cepa*), *AcFT4* expression prevents bulb formation and when overexpressed in *Arabidopsis* significantly delays flowering (Lee *et al*., 2013). *BvFT1* in beets (*Beta vulgaris*) represses flowering through repression of *BvFT2*, the functionally conserved homolog of *FT* in *Arabidopsis* (Pin *et al*., 2010). Similarly, chrysanthemums (*Chrysanthemum seticuspe*) also contain an antiflorigenic FT/TFL1 family protein (CsAFT) that acts antagonistically to CsFTL3 (Higuchi *et al*., 2013). *ZmaFT9* is likely the first antiflorigen to be described in a marine angiosperm where clonal propagation maintains meadows. Further study revealing the mechanism of repression will provide insights into the evolution of the strategy by which *Z. marina* and other marine angiosperms regulate flowering.

### *ZmaTFL1a* may play a role in regulating flowering shoot architecture and other processes not related to flowering

While many genes that group within the *FT* or *TFL1* clade have previously described function in regulating flowering, we know that several *PEBP* genes are implicated in developmental processes other than flowering (Wickland & Hanzawa, 2015). In strawberries (*Fragaria vesca*), three *FT* genes collectively impact plant architecture and *FveFT3* has a strong effect on fruit yield (Gaston *et al*., 2021). The *FT* homolog *StSP6A* induced tuberization in potato (*Solanum tuberosum*) (Navarro *et al*., 2011). In addition, other genes and transcription factors can interact with *FT* homologs to regulate various processes, such as *BRANCHED1* (*BRC1*), which interacts with *FT* to suppress floral development in axillary meristems (Niwa *et al*., 2013). *TFL1* homologs are also known to play a role in flowering shoot architecture, with *tfl1 Arabidopsis* plants having short stems with single terminal flowers (Shannon & Meeks-Wagner, 1991; Kobayashi *et al*., 1999). Overexpression of *RCN1* and *RCN2*, the *TFL1* homologs in rice (*Oryza sativa*) prevents stem elongation and affects branching and panicle development in rice plants (Nakagawa *et al*., 2002). *ZmaTFL1a* appears to be either not expressed or expressed at low levels in leaf tissue in vegetative perennial shoots (Fig. 4). In both ecotypes, *ZmaTFL1a* expression increases greatly in rhizome tissue once the floral transition has occurred (Fig. 7 and S7). Paired with the delayed flowering phenotype in *35S:ZmaTFL1a Arabidopsis* (Fig. 2), we hypothesize that *ZmaTFL1a* may play a role in flowering shoot architecture and regulate spathe and inflorescence formation. Further study of *ZmaTFL1a* function and regulation within *Z. marina* populations may yield new insights into increasing seed potential in a population, a key component of increasing population genetic diversity (Kendrick *et al*., 2012) and a strategy currently being used in restoration practices in the eastern United States (Marion & Orth, 2010; Orth *et al*., 2012, 2020).

With further study of the *Z. marina* genome and transcriptome analyses, we expect that other genes related to flowering will be identified and characterized. We also expect that additional study and transcriptomic analyses will reveal insights into the potential functions of the other *ZmaPEBP* candidate genes that were not brought to light by this study due to experimental limitations and lack of genetics and gene-editing tools. Our study focused primarily on flowering onset and therefore did not pursue inquiry in other developmental functions. However, given the wide array of effects the expression of these genes has on various developmental processes in other plant species, we expect that future studies will further elucidate the roles of *PEBP* genes in eelgrass. We also expect that further study of activating and repressing florigens in the context of different eelgrass population and sites on a larger scale will provide more insights to how these genes impact reproduction- and restoration-focused research. Given our environmental conditions, we predict that the timing and regulation of *ZmaPEBP* genes is affected by local environmental conditions and suggest that continued research into the role of *ZmaPEBP* genes in flowering will allow for more strategic and efficient seed-based restoration efforts of eelgrass.

## CONCLUSION

Our study begins to address the long-standing question of why certain eelgrass shoots flower in a given season and why flowering frequency varies across spatial scales and seasonally. There is a breadth of work exploring this question from the population level, often comparing local environmental factors (Phillips *et al*., 1983; Yang *et al*., 2013; Blok *et al*., 2018; von Staats *et al*., 2021; Ruesink *et al*., 2022), but no studies to date approaching this question from a molecular and mechanistic perspective. Rising atmospheric and water temperatures and increasing frequency of extreme climate events predicted as a result of climate change will likely increase the thermal stress on marine and aquatic ecosystems and have impacts on eelgrass population health (Hobday *et al*., 2018; Jager *et al*., 2018; Kendrick *et al*., 2019). With global loss of seagrass populations estimated at 7% annually and rising (Orth *et al*., 2006; Waycott *et al*., 2009) and given the importance of seeds in promoting resiliency in eelgrass populations (Kendrick *et al*., 2012), a key gap has been understanding how climate change and warming events will impact eelgrass reproduction and persistence. Characterization of the interacting genetic by environmental controls of flowering will lead to a better understanding of factors affecting seed production and to improved seed-based restoration strategies. Understanding how local climate and environmental stimuli affect plant reproductive processes may provide insight into how populations will respond to climate change and help inform their restoration and management.

## ACKNOWLEDGEMENTS

We thank A. K. Hempton, W. Albers, C. Nguyen, K. Kranzler, and A. Keely for technical support and assistance. This research was funded by Washington State Department of Natural Resources Interagency Agreements (93-102930 to J. L. R. and T. I.., and 93-106455 to T. I.), U.S. Geological Survey Northwest Climate Adaptation Science Center award G17AC00218 to C. T. N., the University of Washington Biology Department (W. T. Edmonson Award, Lawrence Giles Botanical Field Research Award, and the Kruckeberg-Walker Award to C. T. N., and the Frye-Hotson Rigg Award to I. C.), and in part by the National Institute of Health (R01GM079712 to T.I.). The views expressed in this article are those of the authors and do not necessarily represent the views or policies of the U.S. Environmental Protection Agency.

## COMPETING INTERESTS

The authors declare no competing interest.

## AUTHOR CONTRIBUTIONS

C. T. N, C. D., J. L. R., and T. I. designed the research. C. T. N., I. C., A. F.-S., and B. A. B. O. performed research. C. T. N., I. C., A. F.-S., B. A. B. O., K. A. N., V. D. S., J.E.K., and J. L. R. analyzed, collected, and/or interpreted data. C. T. N, I. C., J. L. R., and T. I. obtained funding. C. T. N., B. A. B. O., K. A. N., J. L. R., and T. I. wrote the manuscript, together with all authors’ contribution.

## DATA AVAILABILITY

All alignments, scripts, population metadata, and genotypes (filtered and unfiltered) will be available on Dryad at the following link upon acceptance. https://doi.org/10.5061/dryad.sbcc2frh0

## Notes

### Competing Interest Statement

The authors have declared no competing interest.

