## Supplementary Material for "Florigen and antiflorigen gene expression correlates with reproductive state in a marine angiosperm, *Zostera marina*"

New Phytologist

***New Phytologist* Supporting Information**

Article acceptance date: Click here to enter a date.

The following Supporting Information is available for this article:

**Figure S1:** Amino acid sequence alignment of ZmaFT, ZmaTFL1, and ZmaMFT proteins with *Arabidopsis* FT and TFL1 and *Oryza sativa* Hd3a (FT ortholog).
**Figure S2:** Expression levels of *ZmaPEBP* genes in $T_1$ overexpression lines under a *35S* promoter.
**Figure S3:** Characterization of *ZmaFT2, ZmaFT4, ZmaFT9* and *ZmaTFL1a Arabidopsis* overexpressor.
**Figure S4:** Population structure and genetic relationships of eelgrass populations used in our study.
**Figure S5:** Relative expression levels of *ZmaFT2*, *ZmaFT4*, *ZmaFT9*, and *ZmaTFL1a* across different leaf tissues.
**Figure S6:** Representative *Z. marina* annual shoots from Stackpole Annual in April, May, June, and July timepoints.
**Figure S7**: Expression of *ZmaFT4* and *ZmaTFL1A* genes in leaf and rhizome tissue across the 2023 growing season at ST-ANN and NP-ANN sites.
**Figure S8:** Expression of *ZmaFT2*, *ZmaFT4*, *ZmaFT9* and *ZmaTFL1A* genes in leaf and rhizome tissue across the 2023 growing season at YQ-ANN site.
**Table S1:** Accession and GenBank numbers for DNA sequences used in phylogenic analysis.
**Table S2:** qPCR primers used in this study.
**Table S3:** Primers used for pENTR-cloning in this study.
**Table S4:** GPS coordinates and tidal elevation of study sites within Willapa Bay.
**Table S5:** Filtering steps applied to the sequenced reads.
**Table S6:** Pairwise genetic divergence of populations sites in our study.
**Table S7:** Three-way ANOVA results of linear model testing gene expression in relation to three factors (Site, Life stage, Tissue Type)
**Table S8**: Two-way ANOVA results of linear model testing gene expression in relation to two factors (Site, Leaf tissue)
**Table S9:** Two-way ANOVA results gene expression in relation to time in time-series at Stackpole Annual (ST-ANN) and Nahcotta Port Annual (NP-ANN) sites.
**Table S10:** Statistically significant groupings of time-points at Stackpole Annual (ST-ANN) and Nahcotta Port Annual (NP-ANN) site based on post-hoc Tukey tests.
**Table S11**: One-way ANOVA results gene expression in relation to time in time-series at Yaquina Bay Annual (YQ-ANN) site.
**Table S12:** Statistically significant groupings of time-points at Yaquina Bay Annual (ST-ANN) and Nahcotta Port Annual (NP-ANN) site based on post-hoc Tukey tests.
**References**

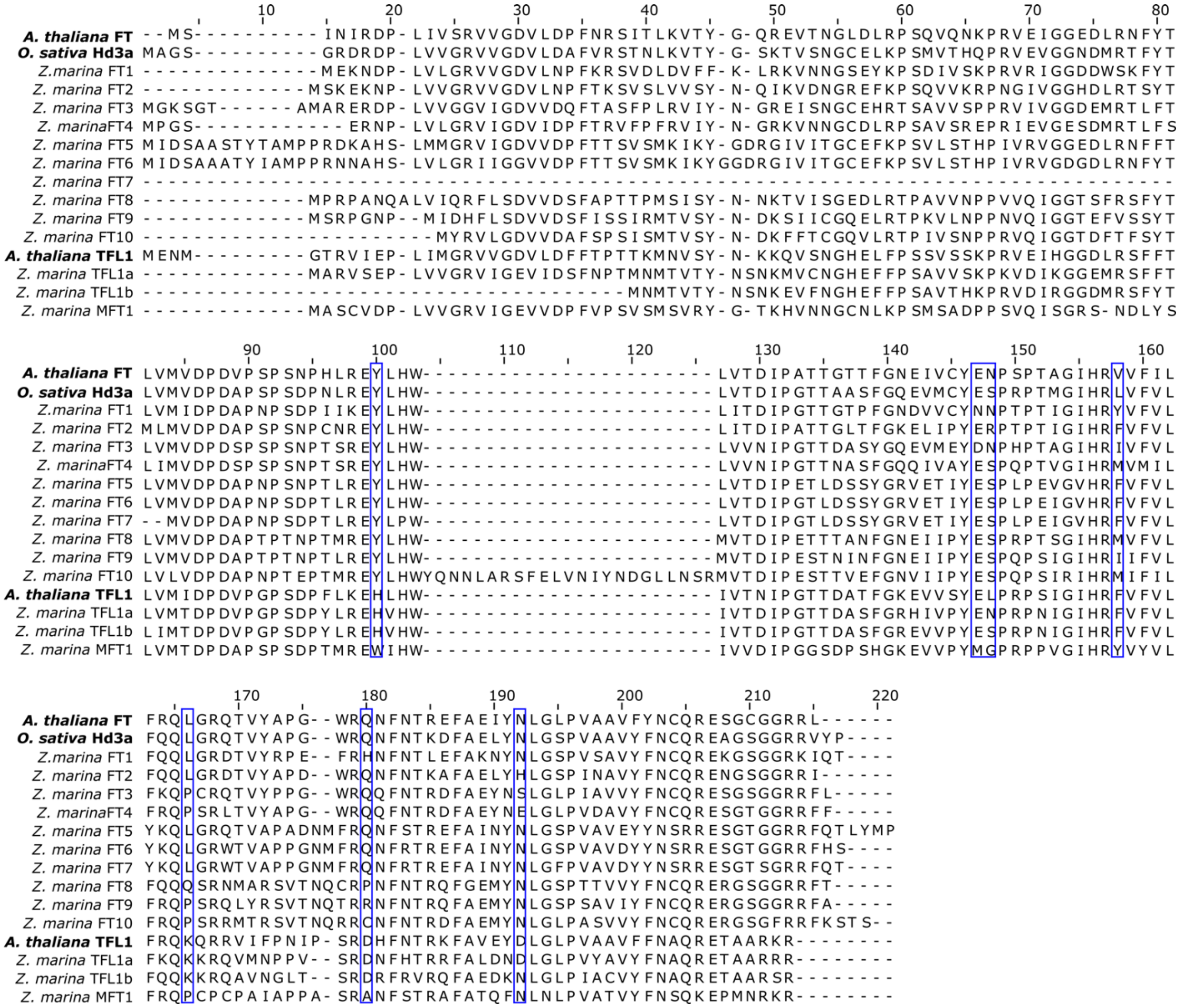

**Figure S1:** Amino acid sequence alignment of ZmaFT, ZmaTFL1, and ZmaMFT proteins with *Arabidopsis* FT and TFL1 and *Oryza sativa* Hd3a (FT ortholog). Blue boxes highlight residues important for function of *Arabidopsis* FT as flowering activator (Y85, E109, N110, V120, L128, Q140, and N152).

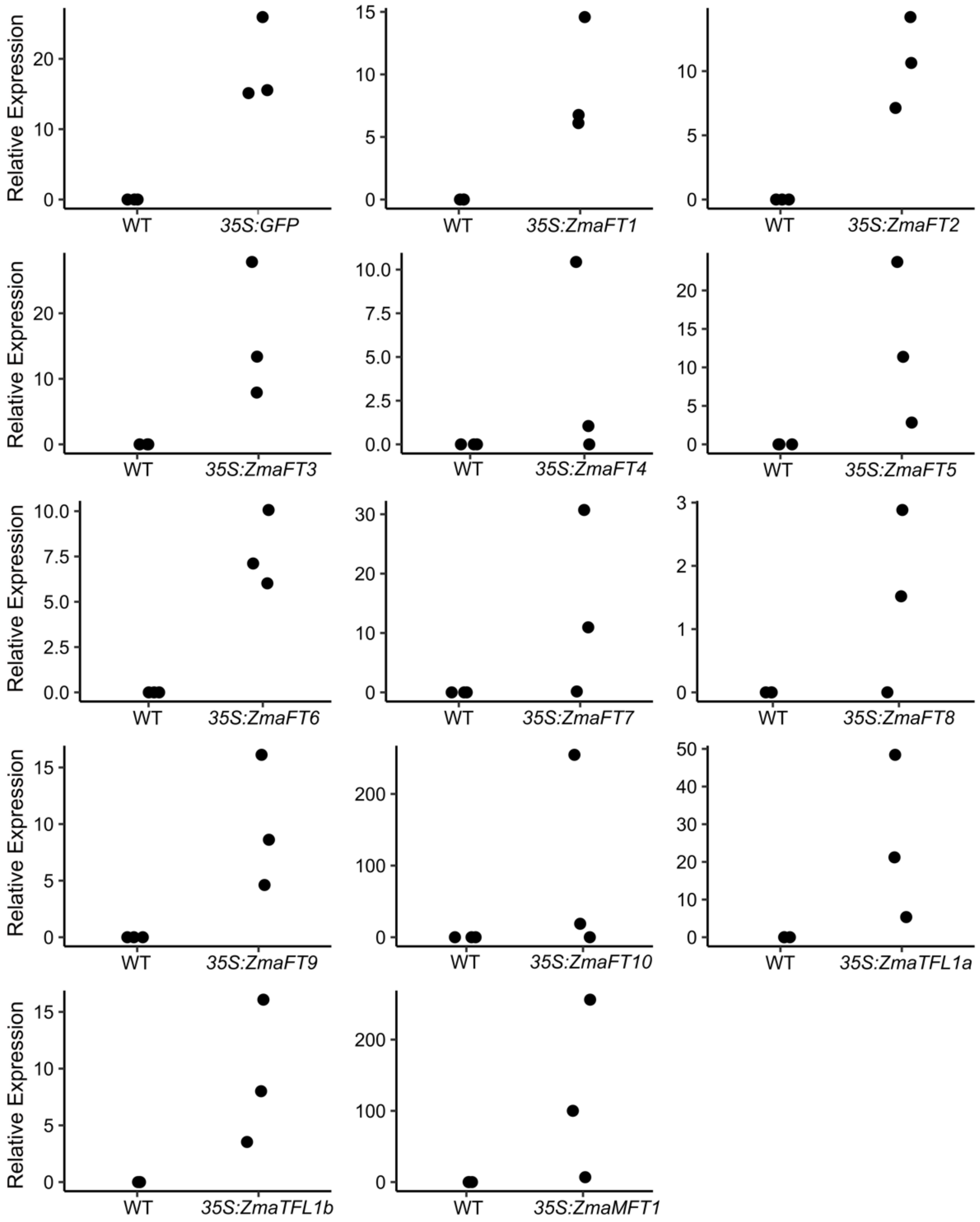


**Figure S2:** Expression levels of *ZmaPEBP* genes in $T_1$ overexpression lines under a *35S* promoter. All lines are in Col-0 background. Each point represents a biological replicate of 4-6 individual $T_1$ plants pooled per sample.

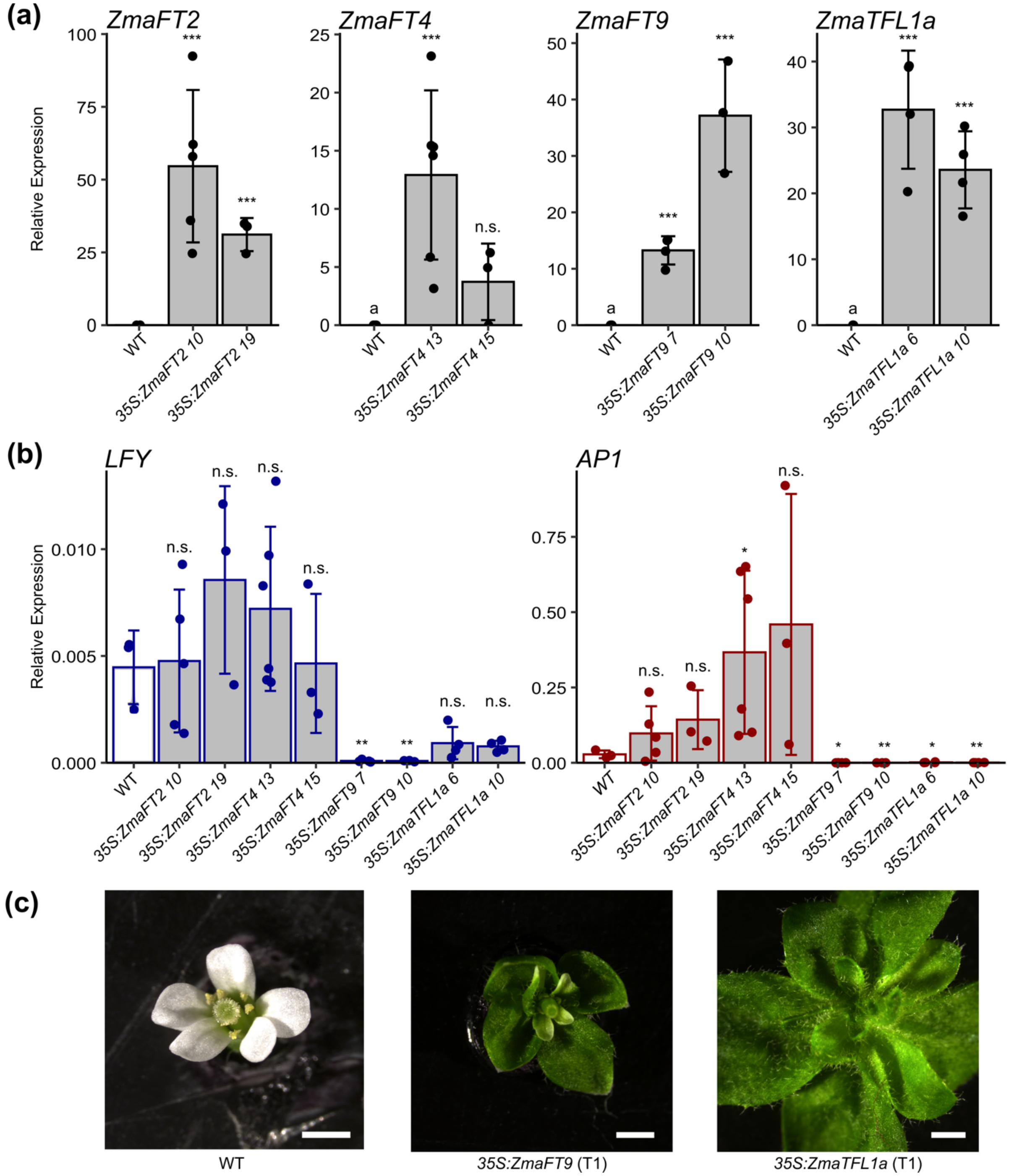


**Figure S3:** Characterization of *ZmaFT2, ZmaFT4, ZmaFT9* and *ZmaTFL1a Arabidopsis* overexpressor. **(a)** Expression of *ZmaFT2, ZmaFT4, ZmaFT9* and *ZmaTFL1a* genes in homozygous $T_3$ overexpression lines under the *35S* promoter. All lines are in Col-0 background. Numbers on x-axis represent different inbred lines. Asterisks indicate significance based on t-test comparing to wild type (WT) with Bonferroni correction with $\log_{10}$ transformed values (n.s.: not significant, ***: $p < 0.0005$). **(b)** Expression of *LFY* and *AP1* in homozygous $T_3$ overexpression lines. Asterisks indicate significance based on t-test comparing to wild type (WT) with Bonferroni correction with $\log_{10}$ transformed values (n.s.: not significant, *: $p < 0.006$, **: $p <$

0.001). **(c)** We observed altered flower morphology in lines overexpressing *35S:ZmaFT9* and *35S:ZmaTFL1* compared to WT (Col-0) in $T_1$ generation. Scale bar 1mm.

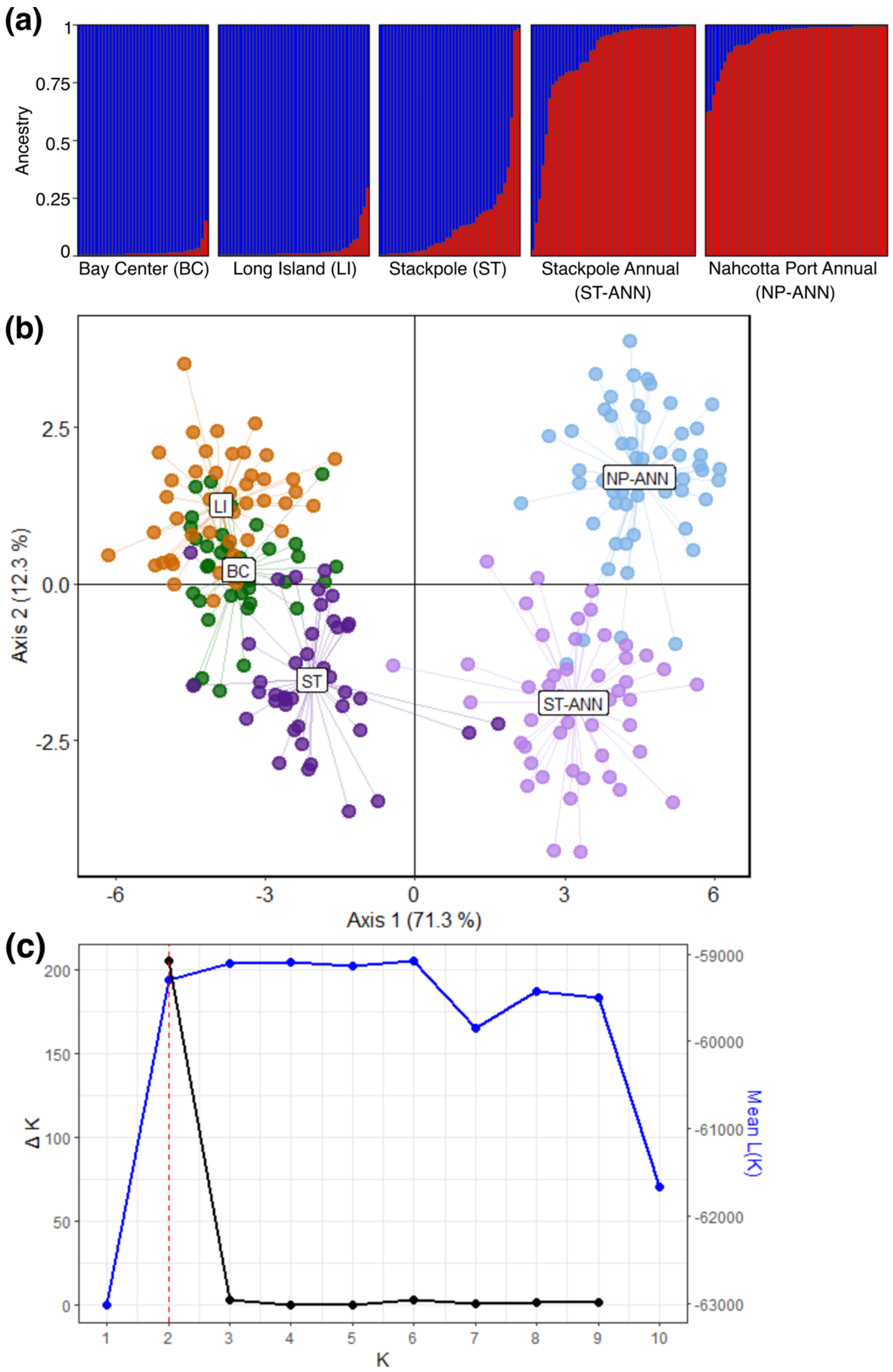


**Figure S4:** Population structure and genetic relationships between eelgrass populations used in our study. **(a)** Population structure inferred using ADMIXTURE analysis implemented in

STRUCTURE v2.1.1.5 (admixture model with correlated allele frequencies, with a burn-in period of 10,000 iterations and 100,000 MCMC repetitions). Each individual plant is represented by a stacked column (x axis), giving the probability of assignment to two ancestral populations (K = 2, Fig. 2c) **(b)** Scatterplot describing genetic relationships among individuals using the first two principal components of DAPC (variance explained in parentheses). Stratified cross-validation (90% of observations used for the training dataset) was performed to identify the optimal numbers of principal components to retain for the DAPC analysis. Data points represent individuals, colors indicate source populations. **(c)** Relationship between K (the number of ancestral populations) and the Mean likelihood of the data $L$(K) (blue) and ΔK (black) values for STRUCTURE analyses. The dashed red line indicates the suggested optimal K value based on the analyses.

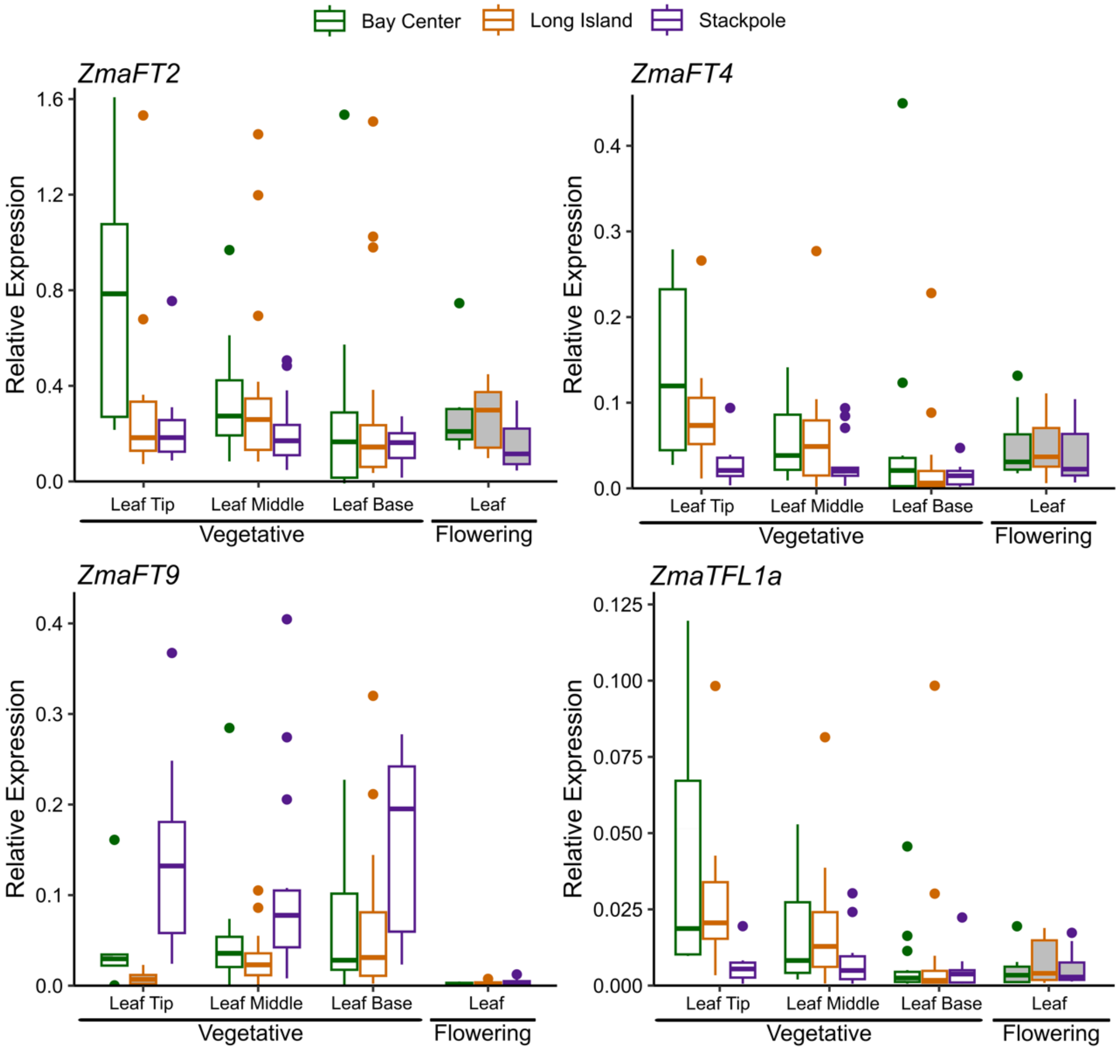


**Figure S5:** Relative expression levels of *ZmaFT2*, *ZmaFT4*, *ZmaFT9*, and *ZmaTFL1a* across different leaf tissues. Results from leaves on vegetative and flowering shoots are shown with white and grey boxes, respectively. Expression in tissues was measured in samples from 3 sites. All expression values are relative to 3 reference genes (*CYP2*, *ELF4A*, and *RPL28*). Median is indicated by center line in box, upper and lower quartiles by box boundaries, and highest and lowest values within two interquartile ranges by whiskers, n≥5.

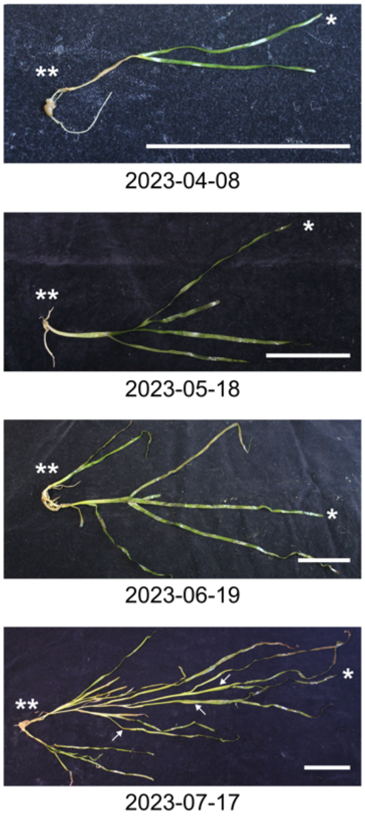


**Figure S6:** Representative pictures of *Z. marina* annual shoots from Stackpole Annual in April, May, June, and July timepoints. Asterisks indicate leaf (*) and rhizome and root (**) tissue sampled. Inflorescences indicated by arrows. Scale bar 5cm.

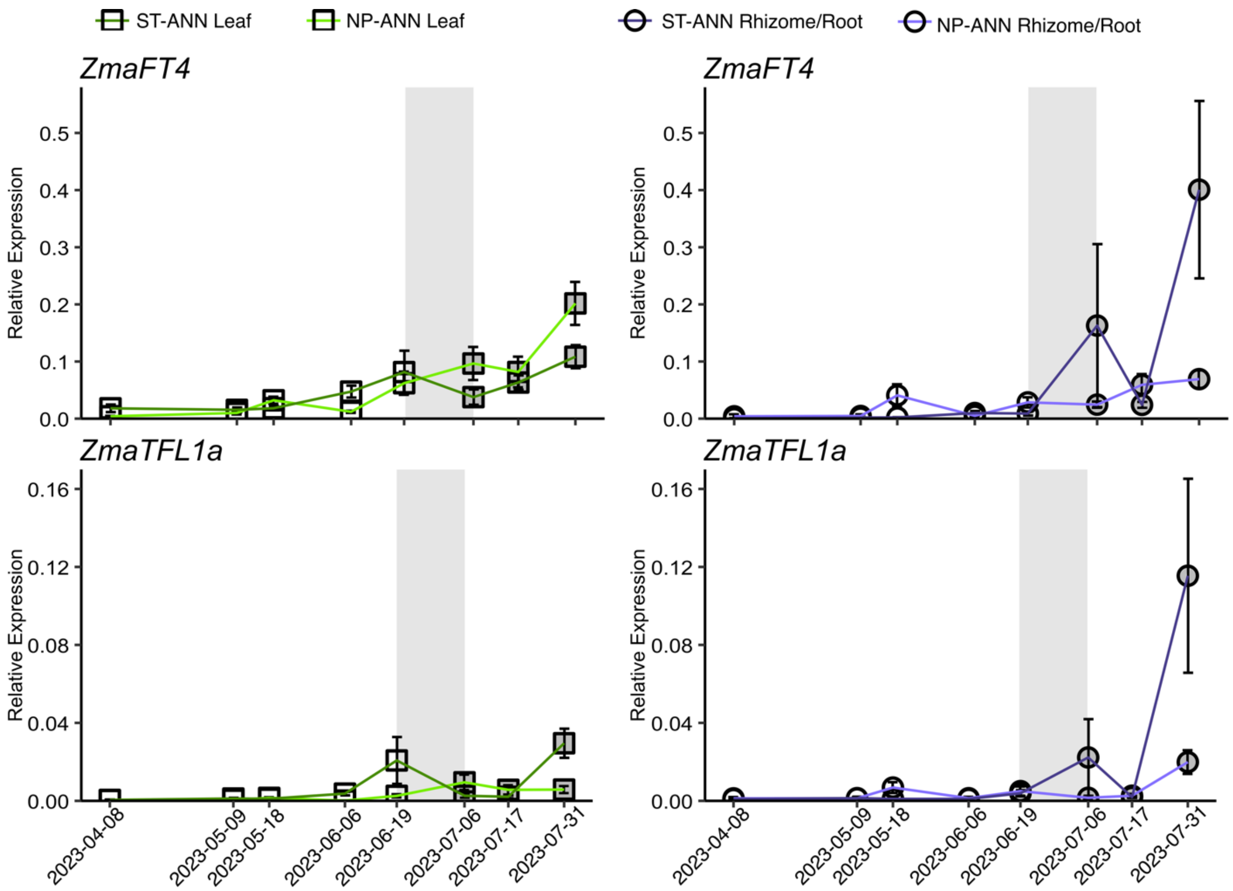


**Figure S7:** Expression of *ZmaFT4* and *ZmaTFL1A* genes in leaf and rhizome tissue across the 2023 growing season at ST-ANN and NP-ANN sites. Grey shaded region in each panel indicates when flowering transition occurred during growing season (April-July). Leaf (square) and rhizome/root (circle) values with lighter green and purple lines indicate NP-ANN site, while darker green and purple lines indicate ST-ANN site. Time points where shoots are vegetative shown in white and time points where shoots had flowered are filled grey. All expression values are relative to 3 reference genes (*CYP2*, *ELF4A*, and *RPL28*). Plot point represents mean, and error bars are standard error. Statistically significant groupings based on post-hoc Tukey HSD of $\log_{10}$ transformed values are listed in Table S10.

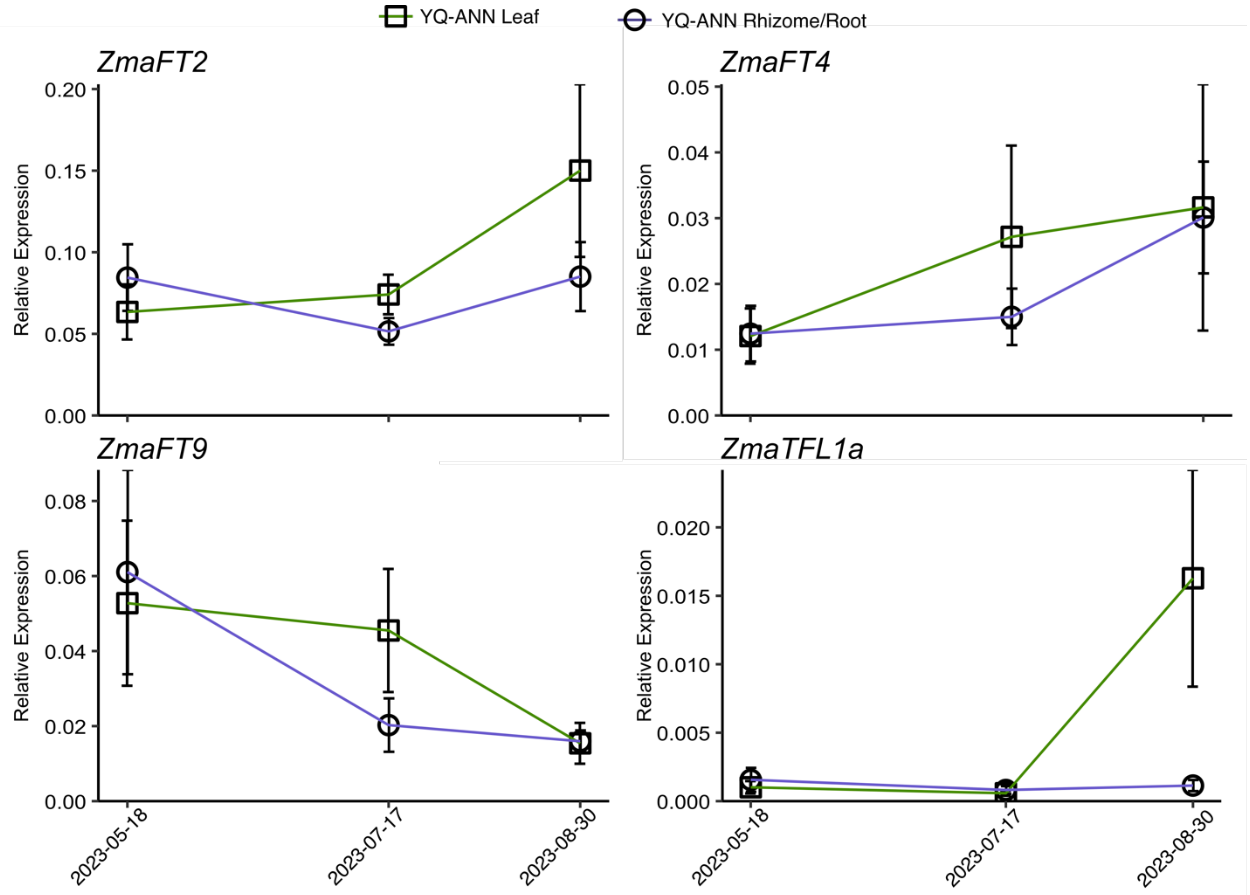


**Figure S8:** Expression of *ZmaFT2*, *ZmaFT4*, *ZmaFT9* and *ZmaTFL1A* genes in leaf and rhizome tissue across the 2023 growing season at YQ-ANN site. Relative expression levels across leaf (square) and rhizome and root (circle) shown throughout the growing season (May-August). All expression values are relative to 3 reference genes (*CYP2*, *ELF4A*, and *RPL28*). Plot point represents mean, and error bars are standard error. Statistically significant groupings based on post-hoc Tukey HSD of $\log_{10}$ transformed values are listed in Table S12. n≥3.

New Phytologist

**Table S1:** Accession and GenBank numbers for DNA sequences used in phylogenic analysis. If gene was functionally characterized, function and reference are also included.

| Name | Species | Gene ID | Database | Function | Publication |
|---|---|---|---|---|---|
| *ZmaFT1* | *Zostera marina* | Zosma03g21100.1 | Phytozome | | |
| *ZmaFT2* | *Zostera marina* | Zosma01g14870 | Phytozome | | |
| *ZmaFT3* | *Zostera marina* | Zosma01g23380.1 | Phytozome | | |
| *ZmaFT4* | *Zostera marina* | Zosma04g08400.1 | Phytozome | | |
| *ZmaFT5* | *Zostera marina* | Zosma06g26410.1 | Phytozome | | |
| *ZmaFT6* | *Zostera marina* | Zosma01g37360.1 | Phytozome | | |
| *ZmaFT7* | *Zostera marina* | Zosma01g37330 | Phytozome | | |
| *ZmaFT8* | *Zostera marina* | Zosma05g25830.1 | Phytozome | | |
| *ZmaFT9* | *Zostera marina* | Zosma01g13540.1 | Phytozome | | |
| *ZmaFT10* | *Zostera marina* | Zosma06g06360.1 | Phytozome | | |
| *ZmaMFT1* | *Zostera marina* | Zosma06g00480.1 | Phytozome | | |
| *ZmaTFL1a* | *Zostera marina* | Zosma04g17290.1 | Phytozome | | |
| *ZmaTFL1b* | *Zostera marina* | Zosma04g24270 | Phytozome | | |
| *ZmuMFT1* | *Zostera muelleri* | | Local (SRR1714574) | | (Lee *et al.*, 2016) |
| *ZmuTFL1b* | *Zostera muelleri* | | Local (SRR1714574) | | (Lee *et al.*, 2016) |
| *ZmuTFL1c* | *Zostera muelleri* | | Local (SRR1714574) | | (Lee *et al.*, 2016) |
| *ZmuTFL1a* | *Zostera muelleri* | | Local (SRR1714574) | | (Lee *et al.*, 2016) |
| *ZmuFT4* | *Zostera muelleri* | | Local (SRR1714574) | | (Lee *et al.*, 2016) |
| *ZmuFT3* | *Zostera muelleri* | | Local (SRR1714574) | | (Lee *et al.*, 2016) |
| *ZmuFT2b* | *Zostera muelleri* | | Local (SRR1714574) | | (Lee *et al.*, 2016) |
| *ZmuFT2a* | *Zostera muelleri* | | Local (SRR1714574) | | (Lee *et al.*, 2016) |
| *ZmuFT1b* | *Zostera muelleri* | | Local (SRR1714574) | | (Lee *et al.*, 2016) |
| *ZmuFT1a* | *Zostera muelleri* | | Local (SRR1714574) | | (Lee *et al.*, 2016) |
| *ZmuFT5a* | *Zostera muelleri* | | Local (SRR1714574) | | (Lee *et al.*, 2016) |
| *ZmuFT5b* | *Zostera muelleri* | | Local (SRR1714574) | | (Lee *et al.*, 2016) |
| *ZmuFT8a* | *Zostera muelleri* | | Local (SRR1714574) | | (Lee *et al.*, 2016) |
| *ZmuFT8b* | *Zostera muelleri* | | Local (SRR1714574) | | (Lee *et al.*, 2016) |
| *ZmuFT10a* | *Zostera muelleri* | | Local (SRR1714574) | | (Lee *et al.*, 2016) |
| *ZmuFT10b* | *Zostera muelleri* | | Local (SRR1714574) | | (Lee *et al.*, 2016) |
| *AtATC* | *Arabidopsis thaliana* | AT2G27550 | TAIR | Repressor | (Mimida *et al.*, 2001) |
| *AtBFT* | *Arabidopsis thaliana* | AT5G62040 | TAIR | Repressor | (Yoo *et al.*, 2010) |
| *AtFT* | *Arabidopsis thaliana* | AT1G65480 | TAIR | Activator | (Kardailsky *et al.*, 1999) |
| *AtMFT* | *Arabidopsis thaliana* | AT1G18100 | TAIR | Activator | (Yoo *et al.*, 2004) |
| *AtTFL1* | *Arabidopsis thaliana* | AT5G03840 | TAIR | Repressor | (Kardailsky *et al.*, 1999; Kobayashi *et al.*, 1999) |
| *AtTSF* | *Arabidopsis thaliana* | AT4G20370 | TAIR | Activator | (Yamaguchi *et al.*, 2005) |
| *BdMFTA* | *Brachypodium distachyon* | Bradi2g01020 | Phytozome | | |

| | | | | | |
|---|---|---|---|---|---|
| *BdMFTBa* | *Brachypodium distachyon* | Bradi2g27860 | Phytozome | | |
| *BdMFTBb* | *Brachypodium distachyon* | Bradi1g42510 | Phytozome | | |
| *BdTCB2* | *Brachypodium distachyon* | Bradi3g44860 | Phytozome | | |
| *BdTCB1* | *Brachypodium distachyon* | Bradi4g42400 | Phytozome | | |
| *BdTCB3* | *Brachypodium distachyon* | Bradi5g09270 | Phytozome | | |
| *BdFT1* | *Brachypodium distachyon* | Bradi1g48830 | Phytozome | Activator | (Lv *et al.*, 2014) |
| *BdFT2* | *Brachypodium distachyon* | Bradi2g07070 | Phytozome | Activator | (Shaw *et al.*, 2019) |
| *BdFT3* | *Brachypodium distachyon* | Bradi2g19670 | Phytozome | | |
| *BdFT13* | *Brachypodium distachyon* | Bradi2g49795 | Phytozome | Activator (SD), Repressor (LD) | (Qin *et al.*, 2019) |
| *BdFT4* | *Brachypodium distachyon* | Bradi1g38150 | Phytozome | | |
| *BdFT6* | *Brachypodium distachyon* | Bradi3g08890 | Phytozome | | |
| *BdFT7a* | *Brachypodium distachyon* | Bradi4g39730 | Phytozome | | |
| *BdFT7b* | *Brachypodium distachyon* | Bradi4g39760 | Phytozome | | |
| *BdFT7c* | *Brachypodium distachyon* | Bradi4g39750 | Phytozome | | |
| *BdFT9* | *Brachypodium distachyon* | Bradi5g14010 | Phytozome | | |
| *BdFT12* | *Brachypodium distachyon* | Bradi3g48036 | Phytozome | | |
| *BdFT10* | *Brachypodium distachyon* | Bradi4g35040 | Phytozome | | |
| *AqFT1* | *Aquilegia coerulea* | Aqcoe2G432300 | Phytozome | | (Sharma *et al.*, 2019) |
| *AqFT2* | *Aquilegia coerulea* | Aqcoe4G263600 | Phytozome | | (Sharma *et al.*, 2019) |
| *AqFT3* | *Aquilegia coerulea* | Aqcoe4G257600 | Phytozome | | (Sharma *et al.*, 2019) |
| *AqMFT* | *Aquilegia coerulea* | Aqcoe3G016200 | Phytozome | | |
| *AqTFL1a* | *Aquilegia coerulea* | Aqcoe2G140500 | Phytozome | | |
| *AqTFL1b* | *Aquilegia coerulea* | Aqcoe1G447800 | Phytozome | | |
| *AqTFL1c* | *Aquilegia coerulea* | Aqcoe1G447700 | Phytozome | | |
| *LaFTL1* | *Lemna aequinoctialis* | LC625473 | GenBank | Activator | (Yoshida *et al.*, 2021) |
| *LaFTL2* | *Lemna aequinoctialis* | LC625474 | GenBank | Repressor | (Yoshida *et al.*, 2021) |
| *SrMFT* | *Symplocarpus renifolius* | LC030438 | GenBank | | (Ito-Inaba *et al.*, 2016) |
| *SrFT* | *Symplocarpus renifolius* | LC030437 | GenBank | Activator | (Ito-Inaba *et al.*, 2016) |
| *SpFT* | *Spirodela polyrhiza* | Spipo19G0001700 | Phytozome | | |
| *CsFTL1* | *Chrysanthemum seticuspe* | AB679270 | GenBank | | |
| *CsFTL2* | *Chrysanthemum seticuspe* | AB679271 | GenBank | | |
| *CsFTL3* | *Chrysanthemum seticuspe* | AB679272 | GenBank | Activator | (Oda *et al.*, 2012) |
| *CsTFL1* | *Chrysanthemum seticuspe* | AB839767 | GenBank | Repressor | (Higuchi & Hisamatsu, 2015) |

| | | | | | |
|---|---|---|---|---|---|
| | *Chrysanthemum seticuspe* | AB770478 | GenBank | | |
| *CsAFT* | *Chrysanthemum seticuspe* | AB839766 | GenBank | Repressor | (Higuchi *et al.*, 2013) |
| *Hd3a/OsFTL2* | *Oryza sativa* | Os06g06320 | Phytozome | Activator | (Kojima *et al.*, 2002) |
| *RFT1/OsFTL3* | *Oryza sativa* | Os06g06300 | Phytozome | Activator | (Kojima *et al.*, 2002) |
| *OsMFT2* | *Oryza sativa* | Os01g02120 | Phytozome | | |
| *OsMFT1* | *Oryza sativa* | Os06g30370 | Phytozome | | |
| *OsRCN1* | *Oryza sativa* | Os11g05470 | Phytozome | Repressor | (Nakagawa *et al.*, 2002) |
| *OsRCN3* | *Oryza sativa* | Os12g05590 | Phytozome | | |
| *OsRCN4* | *Oryza sativa* | Os04g33570 | Phytozome | | |
| *OsRCN2* | *Oryza sativa* | Os02g32950 | Phytozome | Repressor | (Nakagawa *et al.*, 2002) |
| *OsFTL11* | *Oryza sativa* | Os11g18870 | Phytozome | | |
| *OsFTL5* | *Oryza sativa* | Os02g39064 | Phytozome | | |
| *OsFTL6* | *Oryza sativa* | Os04g41130 | Phytozome | | |
| *OsFTL7* | *Oryza sativa* | Os12g13030 | Phytozome | | |
| *OsFTL1* | *Oryza sativa* | Os01g11940 | Phytozome | | |
| *OsFTL8* | *Oryza sativa* | Os01g10590 | Phytozome | | |
| *OsFTL13* | *Oryza sativa* | Os02g13830 | Phytozome | | |
| *OsFTL12* | *Oryza sativa* | Os06g35940 | Phytozome | | |
| *OsFTL4* | *Oryza sativa* | Os09g33850 | Phytozome | Repressor | (Gu *et al.*, 2022) |
| *OsFTL10* | *Oryza sativa* | Os05g44180 | Phytozome | | |
| *OsFTL9* | *Oryza sativa* | Os01g54490 | Phytozome | | |
| *AcFT1* | *Allium cepa* | KC485348 | GenBank | Activator | (Lee *et al.*, 2013) |
| *AcFT2* | *Allium cepa* | KC485349 | GenBank | Activator | (Lee *et al.*, 2013) |
| *AcFT3* | *Allium cepa* | KC485350 | GenBank | | |
| *AcFT4* | *Allium cepa* | KC485351 | GenBank | Repressor | (Lee *et al.*, 2013) |
| *AcFT5* | *Allium cepa* | KC485352 | GenBank | | |
| *AcFT6* | *Allium cepa* | KC485353 | GenBank | | |

**Table S2:** qPCR primers used in this study.

| Name | Accession No. | Forward 5'-3' | Reverse 5'-3' |
|---|---|---|---|
| *ZmFT1* | Zosma03g21100 | GTTCGGATCGGTGGTGATGA | ATGGTCGGCGTTGGGTTATT |
| *ZmFT2* | Zosma01g14870 | ACACAATGCTCATGGTAGATCCT | AGCACAAACACGAAACGGTG |
| *ZmFT3* | Zosma01g23380 | GCCCAAGCAACCCTACTTCT | ACCGTCTGCCTACATGGTTG |
| *ZmFT4* | Zosma04g08400 | TGCACTGGTTGGTGGTGAA | AGCCGACTAGGTTGACGGAA |
| *ZmFT5* | Zosma06g26410 | CGAGTTGGAGGCGAGGATTT | CCGTCACCAACCAGTGTAGG |
| *ZmFT6* | Zosma01g37360 | TGCACACTCTCTTGTGCTGG | CCACGATCACCACCGTACTT |
| *ZmFT7* | Zosma01g37330 | ACCCTGGTTGGTGACGGATA | CAATCTCCGGCAACGGACTT |
| *ZmFT8* | Zosma05g25830 | CACTGGATGGTGACGGACAT | GCGGCATTGATTCGTGACAG |
| *ZmFT9* | Zosma01g13540 | TTCCTACACACTGGTTATGGTCG | GATGGATGGTTGTGGGCTCT |
| *ZmFT10* | Zosma06g06360 | AGAGTCCACAACCGAGCATC | AACCGCTACCTCTCTCCCTT |
| *ZmTFL1* | Zosma04g17290 | TGTTCACTGGATTGTGACAGA | CCAGTGCAAATCGACGTGTG |
| *ZmTFL2* | Zosma04g24270 | ACACACTGATCATGACTGACCC | ATCAGTTGTGCCGGGAATGT |
| *ZmMFT1* | Zosma06g00480 | TTCCTTACATGGGTCCACGC | AAACGGTAGCAACAGGGAGG |
| *ZmCyp2* | Zosma05g32090 | CACTCCACTACAAGGGATCGAAA | GGACCTGTATGCTTCTTAACGAAGT |
| *Zmelf4a* | Zosma03g02900 | TCTTTCTGCGATGCGAACAG | TGGATGTATCGGCAGAAACG |
| *ZmRPL28* | Zosma04g25970 | TTCCGCACCTAGGGTTTCG | ATATTGGCGCAGCGATTTTG |
| *AtIPP2* | AT3G02780 | GTATGAGTTGCTTCTCCAGCAAAG | GAGGATGGCTGCAACAAGTGT |
| *AtPP2AA3* | AT1G13320 | GCGGTTGTGGAGAACATGATACG | GAACCAAACACAATTCGTTGCTG |
| *AtLFY* | AT5G61850 | ACGCCGTCATTTGCTACTCT | CTTTCTCCGTCTCTGCTGCT |
| *AtAP1* | AT1G69120 | AGTGGGATCAGCAGAACCAAGGCC | TTGCAGTTGTAAACGGGTTCAAGAGTCAG |
| *GFP* | None | TGAGCAAGGGCGAGGAGCTG | TGCAGATGAACTTCAGGGTCAGCT |

**Table S3:** Primers used for pENTR-cloning in this study, designed to be compatible with pENTR-D TOPO cloning protocol (Invitrogen).

| Name | Accession No. | Forward 5'-3' | Reverse 5'-3' |
|---|---|---|---|
| *ZmFT1* | Zosma03g21100 | CACCATGGAAAAGAATGATCCACTTGTTC | AATCAAGTCTGAATCTTTCTACCTCCAGATC |
| *ZmFT2* | Zosma01g14870 | CACCATGTCAAAAGAAAAAAATCCCCTTG | AATCATATCCTCCTTCCTCCAGATCCA |
| *ZmFT3* | Zosma01g23380 | CACCATGGGAAAGAGTGGCA | AACTACAAGAATCGGCGACCAC |
| *ZmFT4* | Zosma04g08400 | CACCATGCCGGGATCGGA | AACTAAAAAAAACGCCTTCCCCCA |
| *ZmFT5* | Zosma06g26410 | CACCATGGGGTTGATGATTG | AACTAGGGCATATATAAAGTCTGAAACCTT |
| *ZmFT6* | Zosma01g37360 | CACCATGGGGTTAATGATTGATTCAG | AAAATCATGTGAACCTACGGCCA |
| *ZmFT7* | Zosma01g37330 | CACCATGGTGGATCCTGATGCGCCAAATC | AATCAAGTCTGAAACCTTCTACCACTAGTA |
| *ZmFT8* | Zosma05g25830 | CACCATGCCGAGACCGGC | AATCATGTGAACCTACGGCCA |
| *ZmFT9* | Zosma01g13540 | CACCATGTCGAGGCCAGG | AATTATGCGAACCTTCGACCACCA |
| *ZmFT10* | Zosma06g06360 | CACCATGTACCGTGTTCTT | AATTAAGATGTCGATTTGAACCTTC |
| *ZmTFL1* | Zosma04g17290 | CACCATGGCACGAGTTTC | AACTAACGTCTTCTAGCTGCAGT |
| *ZmTFL2* | Zosma04g24270 | CACCATGAATATGACTGTTACCTACAACTC | AATTAGCGACTGCGAGCTGCAGTCTC |
| *ZmMFT1* | Zosma06g00480 | CACCATGGCGAGCTGCGT | AATCAAGTCTGAAACCTTCTACCACTAGTA |

**Table S4:** GPS coordinates and tidal elevation of study sites including individuals sampled per site and the number of unique multilocus genotypes (MLG) used in the population genetic analyses (BC, LI, ST, ST-ANN, and NP-ANN). Tidal elevation reported in meters of mean lower low water (MLLW)

| Site | Abbreviation | Individuals Sampled | Total MLGs | Position (°N, °W) | Tidal Elevation (m MLLW) |
|---|---|---|---|---|---|
| Bay Center | BC | 44 | 38 | 46.652, 123.946 | 0.3 |
| Long Island | LI | 47 | 44 | 46.515, 123.972 | 0.3 |
| Stackpole | ST | 41 | 41 | 46.611, 124.038 | 1 ± 0.1 |
| Stackpole Annual | ST-ANN | 48 | 48 | 46.613, 124.034 | 1 ± 0.1 |
| Nahcotta Port Annual | NP-ANN | 53 | 53 | 46.502, 124.029 | 0.7 |
| Yaquina Bay Annual | YQ-ANN | - | - | 44.622, 124.042 | 0.3 |

**Table S5:** Filtering steps applied to the sequenced reads. Rows describe individual steps in the bioinformatic pipeline. Columns describe the number of individuals, single nucleotide polymorphic sites (SNPs), and missing data remaining in the dataset after the application of each step.

| Step / Filter | Individuals | SNPs | Missing Data |
|---|---|---|---|
| Single Nucleotide Variant calling with ref_map.pl in STACKS (Rochette & Catchen, 2017) | 243 | 10711 | 64.66% |
| Genotype depth ≥ 10 | 243 | 10711 | 76.68% |
| Genotype Allelic Balance (the proportion of reads supporting a variant site) 0.2 ≥ 0.8 | 243 | 10711 | 76.95% |
| Loci with Minor Allele Count > 1 (remove fixed loci) | 243 | 3926 | 43.35% |
| Loci with < 20% Missing Genotypes across individuals | 243 | 1361 | 6.07% |
| One locus per rad-tag (locus with highest Minor Allele Frequency) | 243 | 695 | 5.08% |
| Individuals with < 20% Missing Genotypes across all loci | 233 | 695 | 2.85% |
| Loci with Minor Allele Frequencies ≥ 0.05 in at least one subpopulation | 233 | 330 | 3.81% |
| Loci meeting Hardy Weinberg Equilibrium expectations in at least one subpopulation | 233 | 327 | 3.78% |
| A single individual per Identical Unique Multilocus Genotypes (MLG) | 224 | 327 | 3.81% |

**Table S6:** Pairwise genetic distances, $F_{ST}$ (below diagonal), with corresponding 95% confidence intervals (above diagonal) among five eelgrass meadows. Confidence intervals (95%) were generated using bootstrapping with 10,000 permutations. Significant values are in bold.

| | BC | LI | ST | ST-ANN | NP-ANN |
|---|---|---|---|---|---|
| BC | | (-0.001, 0.005) | (0.008, 0.018) | (0.100, 0.151) | (0.112, 0.171) |
| LI | 0.002 | | (0.008, 0.018) | (0.099, 0.150) | (0.113, 0.171) |
| ST | **0.013** | **0.013** | | (0.064, 0.101) | (0.078, 0.128) |
| ST-ANN | **0.126** | **0.124** | **0.082** | | (0.011, 0.022) |
| NP-ANN | **0.141** | **0.142** | **0.102** | **0.016** | |

**Table S7:** Three-way ANOVA results of linear model testing gene expression in relation to three factors: Site was included only as a main effect, Life stage (Vegetative, Veg. vs Flowering, Flo.) and Tissue type (Leaf, Rhizome, Stem) were included as main effects and two-way interaction. Expression values were $log_{10}$ transformed to fit assumptions of model.

| | | *ZmaFT2* | | | *ZmaFT4* | | | *ZmaFT9* | | | *ZmaTFL1* | | |
|---|---|---|---|---|---|---|---|---|---|---|---|---|---|
| | Df | Mean Sq | F value | Pr(>F) | Mean Sq | F value | Pr(>F) | Mean Sq | F value | Pr(>F) | Mean Sq | F value | Pr(>F) |
| Site | 2 | 0.26328658 | 3.76413507 | 0.02458575 | 0.66584705 | 3.52774566 | 0.03092301 | 1.64557038 | 2.56810085 | 0.07881367 | 1.55594831 | 4.94852901 | 0.0078441 |
| Developmental Stage (Veg., Flo.) | 1 | 12.482944 | 178.465184 | 1.02E-30 | 16.0270346 | 84.9133473 | 1.73E-17 | 55.4320461 | 86.50805 | 9.59E-18 | 0.95331658 | 3.03192254 | 0.08293884 |
| Tissue type | 2 | 0.608279 | 8.69639592 | 0.00022665 | 2.31443909 | 12.2622042 | 8.56E-06 | 57.8105563 | 90.2199872 | 7.37E-30 | 3.20249286 | 10.1851898 | 5.71E-05 |
| Life stage x Tissue | 2 | 4.1011432 | 58.6329056 | 2.05E-21 | 3.76189104 | 19.9309959 | 9.99E-09 | 35.9194815 | 56.0564605 | 1.16E-20 | 5.43229632 | 17.2768438 | 9.90E-08 |
| Residuals | 237 | 0.0699461 | | | 0.18874576 | | | 0.64077327 | | | 0.31442643 | | |

**Table S8:** Two-way ANOVA results of linear model testing gene expression in relation to two factors: Site and Leaf tissue type (Leaf Tip, Leaf Middle, and Leaf Base) were included as main effects. Expression values were $log_{10}$ transformed to fit assumptions of model.

| | | *ZmaFT2* | | | *ZmaFT4* | | | *ZmaFT9* | | | *ZmaTFL1* | | |
|---|---|---|---|---|---|---|---|---|---|---|---|---|---|
| | Df | Mean Sq | F value | Pr(>F) | Mean Sq | F value | Pr(>F) | Mean Sq | F value | Pr(>F) | Mean Sq | F value | Pr(>F) |
| Site | 2 | 0.16581393 | 1.28135145 | 0.28166476 | 0.62693743 | 2.25208832 | 0.10988044 | 7.74909151 | 18.0619056 | 1.5614E-07 | 1.85664635 | 3.29559782 | 0.04063748 |
| Leaf tissue type | 2 | 0.80581079 | 6.22702086 | 0.00271987 | 4.33339856 | 15.5664597 | 1.07E-06 | 1.38558484 | 3.22957892 | 4.33E-02 | 6.9162808 | 12.2765867 | 1.50E-05 |
| Residuals | 237 | 0.12940551 | | | 0.27838048 | | | 0.42902957 | | | 0.56337164 | | |

**Table S9:** Two-way ANOVA results gene expression in relation to time in time-series at Stackpole Annual (ST-ANN) and Nahcotta Port Annual (NP-ANN) site. Leaf tissue expression and root tissue expression data were analyzed separately. Site was included as a fixed effect, and date was included as a main effect. Date was treated as a categorical factor to account for potential non-linear change. Expression values were $\log_{10}$ transformed to fit assumptions of analysis.

| *ZmaFT2* | | | | | | | | |
|---|---|---|---|---|---|---|---|---|
| | Leaf | | | | Rhizome/Roots | | | |
| | Df | Mean Sq | F value | Pr(>F) | Df | Mean Sq | F value | Pr(>F) |
| Site | 1 | 1.695826686 | 20.25837067 | 1.36674E-05 | 1 | 0.271766091 | 1.13629943 | 0.288244726 |
| Date | 7 | 2.605744737 | 31.12826516 | 3.9622E-26 | 7 | 3.754810051 | 15.69948813 | 3.78125E-15 |
| Site x Date | 7 | 0.339047039 | 4.050260936 | 4.35E-04 | 7 | 0.34131959 | 1.427114227 | 0.198876154 |
| Residuals | 147 | 0.083709925 | | | 142 | 0.239167673 | NA | NA |
| *ZmaFT4* | | | | | | | | |
| | Leaf | | | | Rhizome/Roots | | | |
| | Df | Mean Sq | F value | Pr(>F) | Df | Mean Sq | F value | Pr(>F) |
| Site | 1 | 0.054492413 | 0.261400521 | 0.609926689 | 1 | 0.266466368 | 0.465806179 | 0.496034738 |
| Date | 7 | 4.161673425 | 19.96357907 | 1.19261E-18 | 7 | 12.71449224 | 22.22602831 | 4.25622E-20 |
| Site x Date | 7 | 0.717665985 | 3.442649188 | 1.93E-03 | 7 | 1.58468478 | 2.770165581 | 0.009938725 |
| Residuals | 147 | 0.208463293 | | | 142 | 0.572054173 | | |
| *ZmaFT9* | | | | | | | | |
| | Leaf | | | | Rhizome/Roots | | | |
| | Df | Mean Sq | F value | Pr(>F) | Df | Mean Sq | F value | Pr(>F) |
| Site | 1 | 1.668719912 | 3.741062939 | 0.055011616 | 1 | 0.010529664 | 0.014225919 | 0.905228078 |
| Date | 7 | 17.94779202 | 40.23672222 | 3.62271E-31 | 7 | 6.370384477 | 8.606596338 | 8.88854E-09 |
| Site x Date | 7 | 1.347263453 | 3.020397454 | 5.39E-03 | 7 | 1.806533119 | 2.440684919 | 0.021603424 |
| Residuals | 147 | 0.446055022 | | | 142 | 0.740174655 | | |
| *ZmaTFL1a* | | | | | | | | |
| | Leaf | | | | Rhizome/Roots | | | |
| | Df | Mean Sq | F value | Pr(>F) | Df | Mean Sq | F value | Pr(>F) |
| Site | 1 | 5.685059062 | 16.58488043 | 7.57573E-05 | 1 | 0.787561165 | 1.191270967 | 0.276921784 |
| Date | 7 | 6.898858101 | 20.12586597 | 9.00869E-19 | 7 | 7.702785696 | 11.65129183 | 1.17536E-11 |
| Site x Date | 7 | 2.561012623 | 7.47117799 | 1.10E-07 | 7 | 1.065945672 | 1.612357475 | 0.13647906 |
| Residuals | 147 | 0.342785653 | | | 142 | 0.661110013 | | |

**Table S10:** Statistically significant groupings of time-points at Stackpole Annual (ST-ANN) and Nahcotta Port Annual (NP-ANN) site based on post-hoc Tukey tests in leaves (a, b, c, d, e, f) and roots (u, v, w, x, y, z). Expression values were $log_{10}$ transformed to fit assumptions of ANOVA.

| Date | 2023-04-08 | 2023-05-09 | 2023-05-18 | 2023-06-06 | 2023-06-19 | 2023-07-06 | 2023-07-17 | 2023-07-31 |
|---|---|---|---|---|---|---|---|---|
| *ZmaFT2* | | | | | | | | |
| | Leaf | | | | | | | |
| ST-ANN | abc | bcde | def | def | fg | cdef | efg | g |
| NP-ANN | a | ab | def | bcd | def | fg | def | fg |
| | Rhizome/Root | | | | | | | |
| ST-ANN | w | wx | wx | wx | wxy | wxy | xy | z |
| NP-ANN | wx | wx | wxy | wx | xy | wxyz | wxy | yz |
| *ZmaFT4* | | | | | | | | |
| | Leaf | | | | | | | |
| ST-ANN | abcde | ab | bcdef | cdefgh | defgh | bcdefg | efgh | gh |
| NP-ANN | a | abc | bcdefg | abcd | cdefgh | fgh | defgh | h |
| | Rhizome/Root | | | | | | | |
| ST-ANN | uv | uvw | uvwx | vwxy | vwxy | wxy | xyz | z |
| NP-ANN | u | uvwx | wxy | uvw | wxyz | wxyz | yz | yz |
| *ZmaFT9* | | | | | | | | |
| | Leaf | | | | | | | |
| ST-ANN | f | f | cdef | cd | cdef | bc | c | a |
| NP-ANN | def | f | cde | ef | def | cdef | bc | ab |
| | Rhizome/Root | | | | | | | |
| ST-ANN | z | yz | xy | xy | x | xy | xy | yz |
| NP-ANN | yz | xy | yz | xy | xy | xy | xy | yz |
| *ZmaTFL1a* | | | | | | | | |
| | Leaf | | | | | | | |
| ST-ANN | ab | bcd | bcd | de | ef | cde | bcd | f |
| NP-ANN | ab | abc | bcd | a | cde | def | de | de |
| | Rhizome/Root | | | | | | | |
| ST-ANN | w | wx | wx | wxy | xyz | wxy | wxy | z |
| NP-ANN | wx | wx | xyz | wx | wxy | wxyz | wx | yz |

**Table S11**: One-way ANOVA results gene expression in relation to time in time-series at Yaquina Bay Annual (YQ-ANN) site. Leaf tissue expression and root tissue expression data were analyzed separately. Date was treated as a categorical factor to account for potential non-linear change. Expression values were $log_{10}$ transformed to fit assumptions of analysis.

| *ZmFT2* | | | | | | | | |
|---|---|---|---|---|---|---|---|---|
| | | Leaf | | | | Rhizome/Roots | | |
| | Df | Mean Sq | F value | Pr(>F) | Df | Mean Sq | F value | Pr(>F) |
| Date | 2 | 0.266965246 | 3.444448843 | 0.05084795 | 2 | 0.078710562 | 1.177556369 | 0.331934457 |
| Residuals | 21 | 0.07750594 | | | 17 | 0.066842288 | | |
| *ZmFT4* | | | | | | | | |
| | | Leaf | | | | Rhizome/Roots | | |
| | Df | Mean Sq | F value | Pr(>F) | Df | Mean Sq | F value | Pr(>F) |
| Date | 2 | 0.327162804 | 0.511083633 | 0.60711645 | 2 | 0.078710562 | 1.177556369 | 0.331934457 |
| Residuals | 21 | 0.640135552 | | | 17 | 0.066842288 | | |
| *ZmFT9* | | | | | | | | |
| | | Leaf | | | | Rhizome/Roots | | |
| | Df | Mean Sq | F value | Pr(>F) | Df | Mean Sq | F value | Pr(>F) |
| Date | 2 | 0.032054783 | 0.020559571 | 0.97967003 | 2 | 0.592119052 | 0.544761873 | 0.589773426 |
| Residuals | 21 | 1.559117319 | | | 17 | 1.086931888 | | |
| *ZmTFL1* | | | | | | | | |
| | | Leaf | | | | Rhizome/Roots | | |
| | Df | Mean Sq | F value | Pr(>F) | Df | Mean Sq | F value | Pr(>F) |
| Date | 2 | 3.062983934 | 6.229422413 | 0.00751515 | 2 | 0.459257127 | 0.770505006 | 0.47827979 |
| Residuals | 21 | 0.491696297 | | | 17 | 0.596046908 | | |

**Table S12:** Statistically significant groupings of time-points at Yaquina Bay Annual (YQ-ANN) site based on post-hoc Tukey tests. Expression values were $log_{10}$ transformed to fit assumptions of ANOVA.

| Date | 2023-05-18 | 2023-07-17 | 2023-08-30 |
|---|---|---|---|
| *ZmaFT2* | | | |
| YQ-ANN Leaf | a | ab | b |
| YQ-ANN Rhizome/Root | x | x | x |
| *ZmaFT4* | | | |
| YQ-ANN Leaf | a | a | a |
| YQ-ANN Rhizome/Root | x | x | x |
| *ZmaFT9* | | | |
| YQ-ANN Leaf | a | a | a |
| YQ-ANN Rhizome/Root | x | x | x |
| *ZmaTFL1a* | | | |
| YQ-ANN Leaf | a | a | b |
| YQ-ANN Rhizome/Root | x | x | x |
